# Diet, Climatic Conditions, and Sex Affect the Mycobiome of Wild Common Marmosets (*Callithrix jacchus*)

**DOI:** 10.1101/2025.07.06.663406

**Authors:** Benjamin J Gombash, Paul A Garber, Peter M Finnegan, Anna C McKenney, Júlio César Bicca-Marques, Maria Fernanda De la Fuente, Filipa Abreu, Antonio Souto, Nicola Schiel, Katherine R Amato, Elizabeth K Mallott

## Abstract

Recent research on the gut mycobiome, or the fungal portion of the gut microbial community, suggests that it interacts with host physiology and impacts host health. However, fundamental questions about how the mycobiome is assembled remain unanswered. It remains unclear whether the fungi detected in the gut are predominantly residents of the gastrointestinal tract or if they are largely transient and pass through. To address this question, we sought to determine whether host factors (e.g., sex) or external environmental factors (e.g., variable climatic conditions and diet) were more strongly correlated with the gut mycobiome of common marmosets (*Callithrix jacchus*), a primate species living in the semi-arid Caatinga biome of northeastern Brazil. A stronger correlation with host factors, would suggest a more resident mycobiome, while a stronger association with external environmental factors suggests a more transient mycobiome. We collected 52 marmoset fecal samples across a 2-year period and DNA metabarcoding was used to assess both the mycobiome and diet of each sample. We used FUNGuild to assign ecological roles to the fungi, which were sorted into resident (e.g. animal pathogens) and transient (e.g. plant pathogens) groups. We found that mycobiome richness and evenness varied by host sex and correlated with the arthropod component of the diet while mycobiome composition varied between wetter and drier periods and correlated with the plant portion of the diet. The fact that external environmental factors were associated with the presence of specific taxa led us to conclude that the mycobiome is largely made up of transient taxa.

**Importance:** Other than causing infectious diseases, the function of the mycobiome is poorly understood. There is evidence that the mycobiome can play a positive role, such as training the host’s immune system, and a negative role, such as causing cancer and Alzheimer’s disease. To fully understand these associations, basic questions about which taxa live in the host and which pass through, must be addressed. We used DNA metabarcoding and a tool that can group fungi based on their ecological roles, to determine whether the mycobiome of Caatinga marmosets was more strongly correlated with host sex or host diet and environment. We found that the marmoset mycobiome was more strongly correlated with diet and environment and most fungi passed through the host rather than lived in the host’s gut.

## Introduction

The gut mycobiome, i.e. the fungal component of the gut microbiome, plays a role in training the host immune system (1), causes infectious diseases (2), interacts with and causes cancer (3–5), and contributes to the development of neurological conditions such as Alzheimer’s disease (6,7). It also interacts with components of the host’s gut microbiome (8,9). Given that the gut mycobiome is more variable within an individual than the bacterial microbiome (10) and is influenced by host phylogeny (8), diet (11), and the external environment (12,13), it is essential to determine which fungi are core members of the gut mycobiome. For example, it remains unclear whether the gut mycobiome is predominantly composed of taxa that reside in the gut or transient taxa that simply pass through the gut.

Identifying resident fungal taxa and distinguishing them from transient taxa is important for understanding the function of the gut mycobiome and how it responds to both the external environment and host factors (14,15). If fungal taxa are predominantly resident, then associations between the mycobiome and host factors may be direct and have important implications for the role of the mycobiome in disease. However, if the gut mycobiome is mostly transient, these associations may be better interpreted as evidence of an external effect (i.e. temperature, rainfall, changes in diet) with spurious fungal associations. For example, if *Fusarium*, a common gut mycobiome taxon in Tibetan macaques (*Macaca thibetana*), is a gut resident, then it may assist its host in digesting cellulose (16). If it is transient, however, it may simply be consumed alongside high cellulose dietary items, like leaves, that *Fusarium* itself is digesting. Taxa found in feces that are not capable of surviving at mammalian body temperatures (17) support the transiency hypothesis. Additionally, fungal taxa detected in feces exhibited major overlap with those found in its host’s external environment (18), and many function as plant pathogens and ectomycorrhizal fungi (i.e. fungi that are in symbiosis with the roots of plants) (13), which suggests that these taxa could be transient.

Host factors are more likely to impact the residential portion of the gut mycobiome because differences in host state, such as maturation, reproductive state, sex, or disease, may result in changes to the intestinal environment, modulating changes in the diversity and/or composition of the resident mycobiome. However, these differences are less likely to alter the transient portion of the gut mycobiome. Conversely, changes in external factors are expected to impact which microbes are introduced from the environment, altering the transient portion of the gut mycobiome while having a minimal or indirect effect on the residential gut mycobiome (15).

Host factors and changes in the within sample diversity and composition of the gut are reported to affect the mycobiome of mammals (9,16,19–22). Eukaryotic assemblages in the gut (19) and mycobiome composition (9) vary with host phylogeny. However, within sample diversity of the mycobiome was found not to vary between species of primates (9). Nonetheless, within a single host species, reproductive state is related to the within sample diversity and composition of the mycobiome. In the case of Tibetan macaques, mothers had a greater Shannon diversity during the weaning period than during the post-weaning (21). In contrast, age-related associations within the gut mycobiome are less clear. Older individuals have been reported to have richer or more complex mycobiomes (19,22), while mixed interactions between age and the mycobiome have also been found. For example, juvenile Tibetan macaques had higher Shannon diversity than middle aged adults, while the Chao index, a measure of richness, did not vary by age. In the study, however, mycobiome composition was found to vary with age (16). In another study, Tibetan macaque mothers and infants showed no differences in within sample diversity metrics during the weaning and post weaning periods, but their mycobiome composition differed during the post weaning period (21). Overall, host sex was not correlated with several metrics of within sample diversity across multiple studies (16,20,22), but may be associated with differences in community composition (16).

The external environment of many mammals are believed to influence the within sample diversity and composition of their mycobiomes (e.g., humans (22), wild and captive Tibetan macaques (13,16,21), wild and captive long-tailed macaques (*Macaca fascicularis*) (23), multiple species of wild bats (11), laboratory mice (*Mus musculus*) (12), and additional taxa of other nonhuman primates, including apes, platyrrhines, cercopithecoids, and lemurs (9,19)). Also, dietary shifts have been proposed to account, at least partially, for changes in the diversity and composition of the mycobiome attributed to external factors, such as season (16), rural and urban environments (22), and captivity (13,23). However, no study has simultaneously assessed the host diet and gut mycobiome of wild animals. Examining how changes in diet affect host mycobiome is needed because the endophytic mycobiome of leaves consumed or contracted by foragers differs in their species-specific fungal communities (24). Similarly, hundreds of species of fungi are known to be arthropod pathogens, and some of these are host specific (25). Therefore, changes in the fungal communities of host diet are expected to result in changes in the composition of the host gut mycobiome.

In this study, we analyze the diet and mycobiome of wild common marmosets (*Callithrix jacchus*) residing in a Caatinga (semi-arid tropical environment) habitat in northeastern Brazil. Specifically, we examine the degree to which the gut mycobiome is largely resident or transient, leveraging ecological information about the fungal taxa detected in feces (26). Taxa from the Animal Pathogen, Endosymbiont, and Animal Endosymbiont fungal guilds (26) can be classified as likely resident (hereafter resident) as they are known to live on or in animal hosts. In contrast, the Plant Pathogen, Endophyte, Epiphyte, Bryophyte Parasite, Ectomycorrhizal, and Plant Parasite guilds (26) are more likely to be transient (hereafter transient) taxa that are unintentionally consumed alongside dietary resources. Examining how these subsets of the mycobiome respond to host and external environmental factors can help us determine whether the mycobiome is largely made up of resident or transient taxa. This system presents an excellent opportunity to investigate the role of a host factor (sex) and external environmental factors (wetter or drier period and diet) play in shaping the gut mycobiome of common marmosets.

Common marmosets are characterized by a reproductive strategy in which generally a single female in each group regularly produces dizygotic twins (27). Gestation is approximately five months, and lactational anestrous commonly results in the breeding female giving birth to two twin litters per year. Group members, principally adult males, assist the breeding female in provisioning, transporting, and caring for the infants (27,28). Mean group size in the Caatinga is six individuals (29). Furthermore, adult female marmosets inhabiting the Caatinga (one of the driest habitats that support non-human primates) are approximately 20% heavier than adult males (29). These traits, which are not typical of primates, provides an opportunity to investigate the role that sex plays in shaping the composition of the gut mycobiome.

At our study site, mean rainfall is 337 mm per year, with no rainfall during some years. However, the timing of rainfall is highly variable across years, with the dry period lasting 5 to 11 months (30). Thus, rather than experiencing a strict dry and wet season, patterns of rainfall during individual months can have a significant effect on resource productivity, as Caatinga adapted plants quickly respond to an increase in precipitation (31). In this regard, common marmosets appear to capture more insect during wetter periods (71% of observed captures), than during drier periods (28%). However, exudates (gums and saps) from a limited number of tree species, comprise the bulk of the common marmoset diet, and are consumed year-round (32). Similarly, temperature varies minimally across the year, with slightly warmer temperatures during drier periods (mean maximum temperature 31.3°C) than during the wetter periods (mean maximum temperature 29.1°C) (32).

Given the importance of even minor differences in rainfall on the availability of insect and plant resources in a Caatinga habitat, we examined how host diet and rainfall, may shape the gut mycobiome of a wild, but provisioned, population of common marmosets (see Methods). To accomplish this, we address the following research questions: 1) Do male and female common marmosets differ in the composition and diversity of their gut mycobiome, and are these differences independent of potential sex-based dietary differences; 2) Does gut mycobiome composition and diversity vary during wetter and drier periods; 3) Are the diversity and taxonomic composition of a marmoset’s diet correlated with its gut mycobiome; and 4) Is the common marmoset mycobiome best described as resident or transient (Figure 1)?

**Figure 1.**
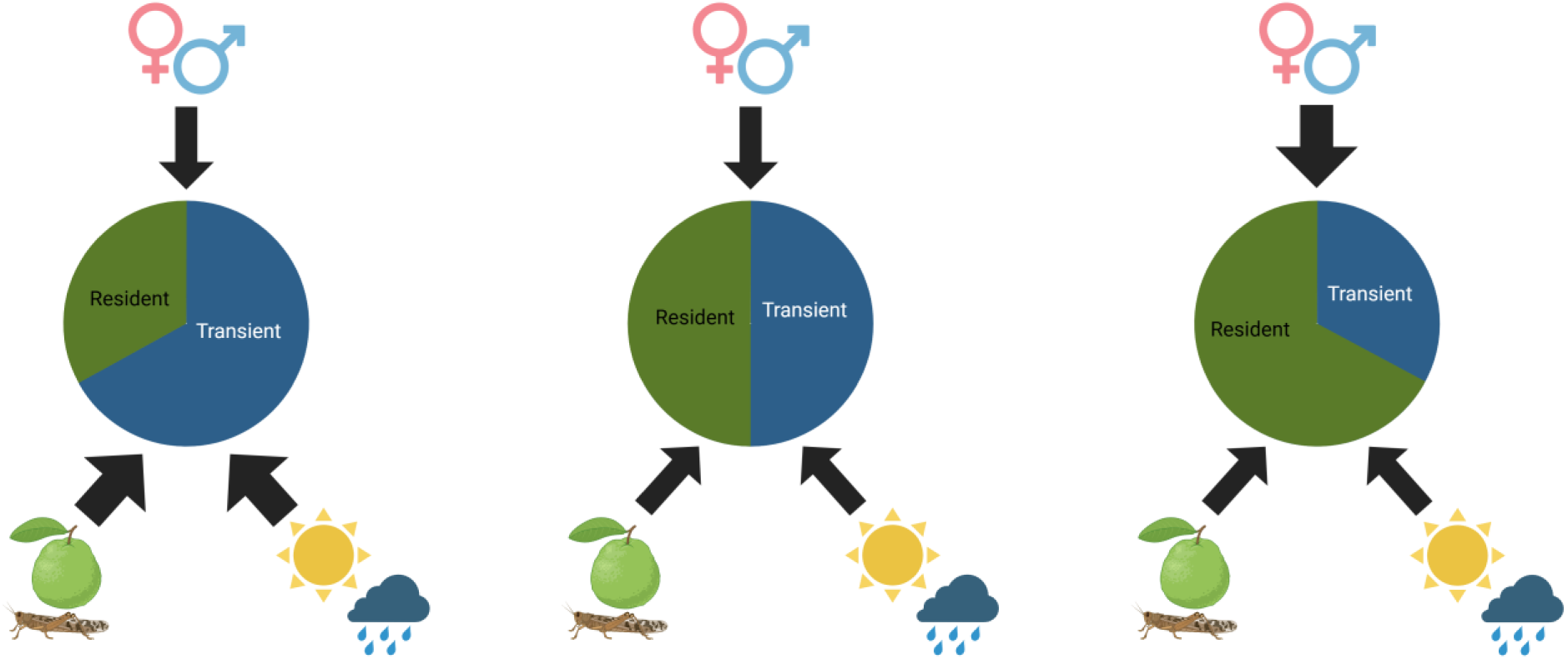
Hypothesized framework showing the expected relative contributions of residential (green) and transient (blue) taxa to the mycobiome under scenarios where environmental factors contribute more than (larger arrows), equal to, or less than (smaller arrows) host factors to the composition of the mycobiome. The specific factors being investigated in this study include sex, diet, and climatic conditions.

## Results

### ITS Sequencing and FUNGuild Results

Sequencing yielded 13,401,893 reads across the 60 samples (mean ± sd = 223,365 ± 174,806 per sample, range = 9,106 to 612,305 per sample). After filtering, pairing, and denoising, we were left with 11,174,377 reads and 2,099 fungal ASVs. Of the ASVs classified as fungi, 787 of which were only classified at the kingdom level, 1,312 ASVs were classified at the phylum or a lower taxonomic level, and 1,188 were classified at the family or below. We assigned 818 ASVs to guilds with FUNGuild and further sorted ASVs into transient or resident categories based on their membership in specific guilds (see Methods). We found 456 transient ASVs and 62 ASVs of resident fungi.

Most of the classified ASVs belonged in the phylum Ascomycota (83.5% of ASVs). Basidiomycota was the next most common phylum (12% of ASVs), followed by Basidiobolomycota (3.7% of ASVs), Chytridiomycota (0.6% of ASVs), and Blastocladiomycota (1 ASV). Ascomycota also made up the largest portion of the reads (56.5% of reads). Reads associated with ASVs that were unidentified to the phylum level were the next largest portion (36.6% of reads). Basidiomycota accounted for 4.6% of reads, Basidiobolomycota made up 1.9% of reads, Chytridiomychota, 0.4% of reads, and Blastocladiomycota only 2 reads.

### Do male and female common marmosets differ in their within sample diversity and composition of the gut mycobiome?

Within sample fungal diversity was significantly higher in male marmosets compared with female marmosets across multiple within sample diversity metrics. Shannon diversity, ASV richness, and Chao1 diversity were all significantly related to the marmoset’s sex (Likelihood ratio tests comparing full mixed-effects models to null models: Figure 2A; Table 1; Figure S1; Tables S1-S3). Simpson’s diversity and Pielou’s evenness differed between sexes (Fixed-effects models: Simpson’s diversity, p-value = 0.008, t-value = 2.814, Figure 2C, Table S4; Pielou’s evenness, p-value = 0.010, t-value = 2.712, Table S5, Figure S2), although the overall models were not significant (Simpson model, p-value = 0.116; Pielou model p-value = 0.133). Sex did not have a significant impact on the mycobiome composition for either distance metric (PERMANOVA, Bray-Curtis: pseudo-F(1) = 1.040, R^2^ = 0.020, p-value = 0.358; Jaccard: pseudo-F(1) = 1.054, R^2^ = 0.021, p-value = 0.141, Figure 2E, Tables S6-7).

**Figure 2.**
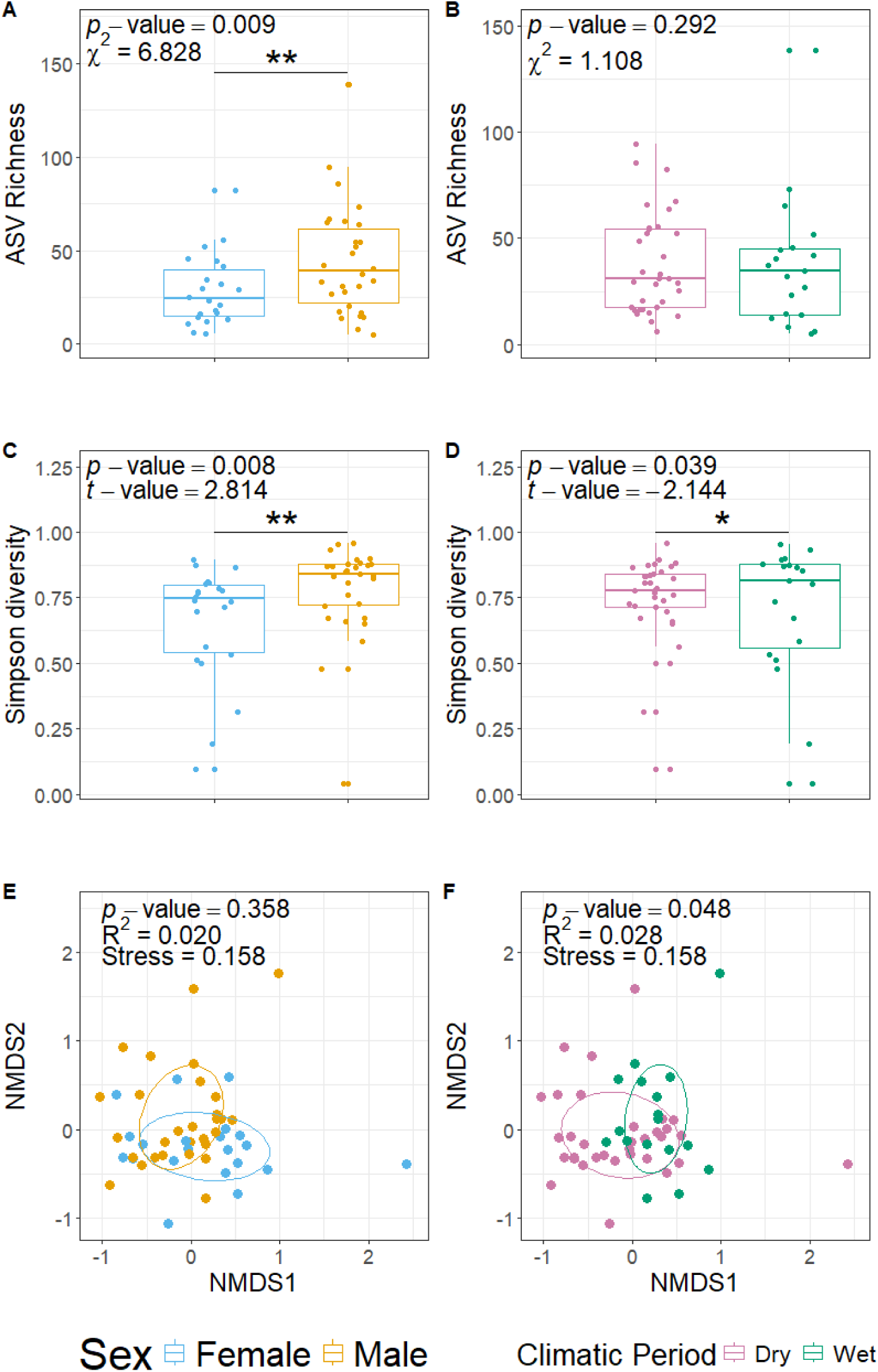
Within sample diversity (A-D) and compositional (E-F) differences among sexes (A, C, and E) and sample collection period (B, D, and F), which are indicated by color. Wald’s test was used to find significant differences between the richness values of sexes and wetter and drier periods (Table S2).

**Table 1.** Results of likelihood ratio tests comparing full mixed-effects models to null models missing the parameter of interest, sex or climatic period, to determine whether they have significant impacts on within sample diversity metrics of the overall mycobiome. Significant models are bolded.

| Diversity Metric | Model | logLik | deviance | Chisq | Df | Pr(>Chisq) |
| --- | --- | --- | --- | --- | --- | --- |
| Shannon | Full Model | -51.472 | 102.94 | 0 | 0 |  |
|  | Null Climatic Period Model | -51.472 | 102.94 |  |  |  |
|  | **Full Model** | **-51.472** | **102.94** | **0** | **0** | **<0.001** |
|  | Null Sex Model | -56.901 | 113.8 |  |  |  |
| Richness | Full Model | -237.86 | 475.73 | 0 | 0 |  |
|  | Null Climatic Period | -237.86 | 475.73 |  |  |  |
|  | **Full Model** | **-237.86** | **475.73** | **6.413** | **1** | **0.011** |
|  | Null Sex Model | -241.07 | 482.14 |  |  |  |
| Chao1 | Full Model | -244.22 | 488.43 | 0 | 0 |  |
|  | Null Climatic Period | -244.22 | 488.43 |  |  |  |
|  | **Full Model** | **-244.22** | **488.43** | **6.409** | **1** | **0.011** |
|  | Null Sex Model | -247.42 | 494.84 |  |  |  |

Seven of the fungal ASVs were differentially abundant between the sexes. A total of 1,840 of the 2,099 identified ASVs remained after removing samples without any information about the sex of the sampled marmoset. Of the seven differentially abundant fungal taxa, six were more abundant in females (*Leptosillia* sp., *Nothophoma* sp., and three unidentified fungi) and two in males (both *Cladosporium* spp.; Figure 3A; Table S8).

**Figure 3.**
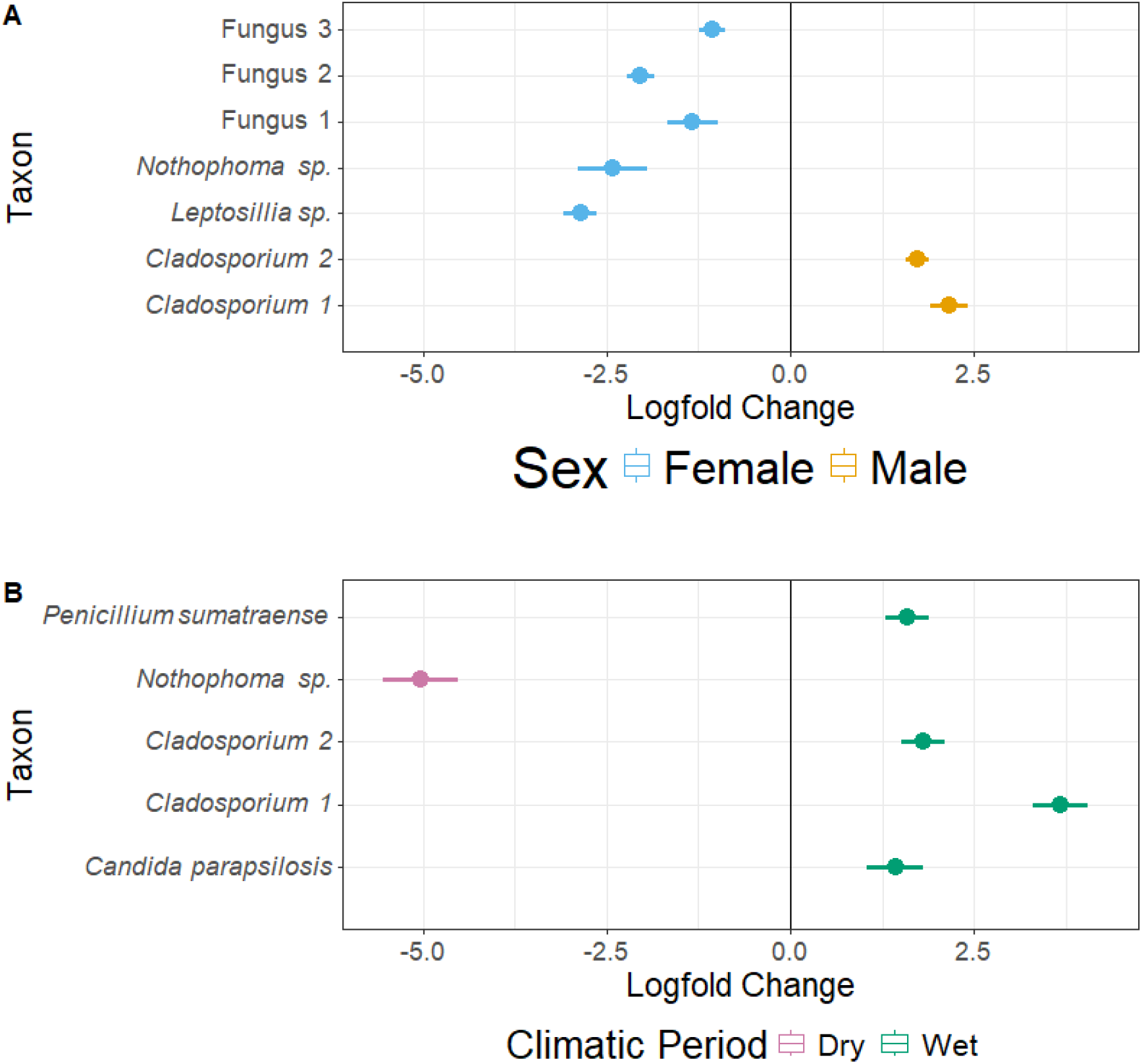
A dot and whisker plot visualizing the statistically significantly differentially abundant taxa indicated by the ANCOMB-BC, which employed a Benjamini-Hochberg procedure, for sex (**A**) and wetter and drier periods (**B**). The dot indicates the logfold change of the specific taxon, with the lines showing the standard error. Colors indicate which category the ASV is associated with.

### Is the marmoset’s sex associated with the within sample diversity of the resident and transient portions of the gut mycobiome?

Sex modulated the ASV richness and the Chao1 diversity of the transient fungi (Likelihood ratio tests comparing full mixed-effects models to null models: ASV richness, χ^2^ = 6.462, p-value = 0.011; Chao1, χ^2^ = 6.180, p-value = 0.013; Table S9), with males having higher richness than females. No other within sample diversity metrics varied by sex (i.e. Shannon diversity, Simpson diversity, and Pielou’s evenness; fixed-effects models; Tables S10-12). Regarding the resident fungi, only Shannon diversity significantly differed by sex, with males having higher diversity (Fixed-effects models, t-value = 2.071, p-value = 0.045, Table S13), although the model itself was not significant (F = 1.046, adjusted R^2^ = 0.012, p-value = 0.431). Other metrics did not vary by sex (Tables S14-17).

### Are the climatic conditions of the sampling period associated with the within sample diversity and composition of the gut mycobiome?

The period of sample collection (wetter vs. drier) was only significantly associated with one metric of within sample diversity. Simpson’s diversity was significantly lower during the drier period (Fixed-effect models, t-value = −2.144, p-value = 0.039, Figure 2, Table S4). Shannon diversity, ASV richness, and Chao1 diversity (Likelihood ratio tests comparing full mixed-effects models to null models, Figure 2B; Table 1; Figure S1; Tables S1-3), and Pielou’s evenness (t-value = −1.693, p = 0.099; Table S5; Figure S2) were not influenced by period. The sampling period significantly impacted the composition of the mycobiome for both distance metrics (PERMANOVA: Bray-Curtis, pseudo-F(1) = 1.475, R^2^ = 0.028, p-value = 0.048; Jaccard, pseudo-F(1) = 1.178, R^2^ = 0.024, p-value = 0.006; Figure 2, Tables S6-7; Figure S5).

Five of the 1,840 ASVs differed in abundance between sample collection periods. A species of *Nothophoma* was more abundant in drier period samples, whereas *Penicillium sumatraense*, *Candida parapsiolosis*, and two species of *Cladosporium* were more abundant in wetter period samples (Figure 3B, Table S18).

### Are the climatic conditions of the sampling period associated with the within sample diversity of the resident and transient portions of the gut mycobiome?

Sampling period did not influence the within sample diversity metrics of transient fungi (i.e. Richness, Chao1, Shannon diversity, Simpson diversity, and Pielou’s evenness; Tables S9-12). The within sample diversity metrics of the resident fungi similarly did not significantly differ by sample collection period (Tables S13-17).

### Does the within sample diversity and composition of a marmoset’s diet, based on DNA in feces, correlate with the diversity of its mycobiome?

There were no significant correlations between within sample diversity metrics of the plant portion of the marmoset diet and the within sample diversity metrics of the overall mycobiome, the transient fungi, or the resident fungi (Table S19). Only ASV richness of the arthropod portion of the diet was significantly positively correlated with the overall mycobiome (rho = 0.287, p-value = 0.039, Figure 4, Table S20). ASV richness for the arthropod portion of the diet was significantly positively correlated with the transient fungi counterpart (rho = 0.293, p-value = 0.035, Figure 4, Table S20), while arthropod Simpson diversity of the diet was the only metric that was significantly positively correlated with its resident fungi counterpart (rho = −0.298, p-value = 0.032, Table S20).

**Figure 4.**
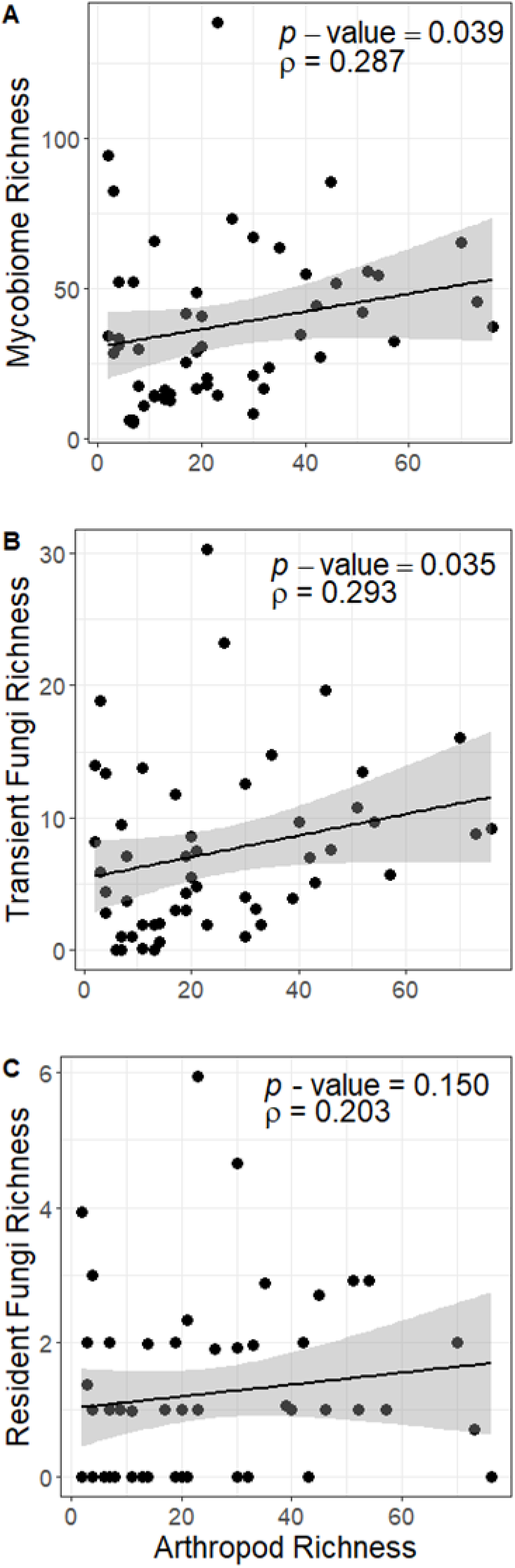
Spearman correlation plots showing correlations between the arthropod richness and the overall mycobiome richness (**A**), transient fungi richness (**B**), and the resident fungi richness (**C**).

The composition of the plant portion of the diet significantly correlated with the overall mycobiome (Mantel r = 0.108, p-value = 0.017) and the resident fungi (Mantel r = 0.248, p-value < 0.001), but not the transient fungi (Mantel r = 0.014, p-value = 0.646). The composition of the arthropod portion of the diet did not significantly correlate with the mycobiome (Mantel r = 0.017, p-value = 0.632), the transient fungi (Mantel r = −0.017, p-value = 0.527), or the resident fungi (Mantel r = 0.079, p-value = 0.056).

We found no significant interactions between 12 fungal and 44 arthropod ASVs (CCREPE: absolute sim score > 0.6, q-value < 0.05, Table S21, Figure S8). We found only one interaction between an unidentified fungus (hereafter Fungus 4) and an *Acacia* species (absolute sim score = 0.630, q-value = 0.023, Table S22, Figure S9) in comparisons between 12 fungal ASVs and 266 plant ASVs. Only six fungal ASVs were included in both CCREPE analyses, including three of the fungi identified by two ANCOM-BC analyses (Fungus 1, *Nothophoma* sp., and *Cladosporium* 1; Figure 3; Tables S8, S18, S21-22). *Candida parapsilosis*, which was associated with the wetter period by ANCOM-BC analysis, appeared in the CCREPE analysis for arthropods and fungi, but did not have any significant pairwise associations (Figure 3; Tables S18 and S21).

### Does the marmoset’s diet vary by sex?

Sex did not correlate with any within sample diversity metric of the arthropod or the plant portions of the diet (Tables S23-28). Sex did not impact the composition of either portion of the diet for either distance metric (Tables S29-32).

## Discussion

In this study we examined the gut mycobiome of Caatinga common marmosets to determine how it correlates with host sex, sample collection period, and host diet. We found that male Caatinga common marmosets had greater within sample diversity of their resident, transient, and overall mycobiome than females. However, the overall mycobiome composition did not vary by sex. Sample collection period was significantly associated only with Simpson diversity of the overall mycobiome, which increased during wetter periods. In contrast, mycobiome composition did vary by period. The within sample diversity of the arthropod component, but not the plant, was positively correlated with that of the overall, transient, and resident mycobiome. We did not find significant associations between the composition of the arthropod diet and the composition of the gut mycobiome (i.e. overall, transient, resident). However, the composition of the mycobiome significantly varied with the composition of the plant portion of the diet. Taken together, the role that external factors, like diet and climatic conditions, play in shaping the composition of the gut mycobiome (i.e. which taxa are present) suggest that the members of the fungal community in the gut of common marmosets are largely passing through the gut alongside the plants and arthropods that they consume; these taxa are not residing and replicating in the host’s gastrointestinal tract. Therefore, we conclude that the common marmoset gut mycobiome is largely transient, as has been proposed for the eukaryome of black and gold howler monkeys, *Alouatta caraya* (33).

### Relationship between sex and mycobiome diversity

We found that male marmosets had significantly greater within sample diversity than females across multiple metrics, which was unexpected given the results of prior studies. In yellow baboons (*Papio cynocephalus*) (20), Tibetan macaques (16), and Seychelles warblers (*Acrocephalus sechellensis*) (34), there were no sex-based differences in gut mycobiome within sample diversity, and, in one study of humans, female individuals had greater within sample diversity than male individuals (35). Differences in humans were proposed to be either caused by hormonal differences or sex-based dietary differences (35). Based on fecal DNA, we did not find any sex-based differences in the marmoset diet, which is consistent with previous studies indicating that common marmosets live in small cohesive social groups, with all or most group members feeding in the same tree at the same time, with low rates of aggression at feeding sites (27,29). Therefore, due to this lack of sex-based dietary differences, hormones may be driving mycobiome differences between sexes, which would mirror interactions between gut bacteria and sex hormones (36). Targeted research in this area is required to fully understand the how host sex interacts with the gut mycobiome.

The fact that we detected differentially abundant taxa between sexes could be interpreted as evidence of a largely resident mycobiome; however, the identities and ecological roles of these taxa suggests that they are transient fungi. The two fungal ASVs that were more abundant in male marmosets were both *Cladosporium* (phylum Ascomycota). While FUNGuild (v1.2; 26) did not assign an ecological role to *Cladosporium*, this genus has been described as a plant pathogen that lives on other fungi and is saprobic (i.e. living on decaying organic matter). It is widely distributed and can be found in “soil, food, … and other organic matters” (37). When *Cladosporium* is acting as a plant pathogen, it can be host specific, but the genus has been found on a wide variety of plant species and plant parts (37). In contrast, six ASVs are more abundant in samples from female marmosets. Only two of these had taxonomic information beyond the kingdom. These ASVs were identified to *Nothophoma* (phylum Ascomycota) and *Leptosillia* (phylum Ascomycota). We considered this *Nothophoma* ASV to be an animal pathogen, plant pathogen, and saprotroph, and it was proposed as a plant associated fungi by Chen et al., (38). There was no information about the function of *Leptosillia* in FUNGuild. However, it has been considered to be a plant endophyte (39). These differential abundances suggest potential microhabitat and/or dietary differences between male and female common marmosets. We did not find any sex-based dietary differences, although metabarcoding data provide a temporal snapshot, confirming a lack of differences would require longitudinal observational data. The sexes may differ in how they use microhabitats (e.g. contact with soil or using different species of plants for travel and rest). If these sex-based differences are related to diet and/or microhabitat use, then they may support the conclusion that the mycobiome is largely transient.

### Relationship between climatic condition and mycobiome diversity

The associations between the within sample diversity metrics, composition, and the sampling period are consistent with a previous study indicating seasonal differences in mycobiome diversity in Tibetan macaques (*M. thibetana*). In that study, mycobiome diversity was greatest during the winter, which is a time of lower temperatures (16). In the case of Tibetan macaques, the Chao index, a measure of richness, showed no significant differences across seasons while the Shannon diversity index, a measure of richness that accounts for the abundance of taxa, was highest in winter and lowest in the summer (16). The authors suggested that the diversity of the mycobiome increased as the diversity of the plant parts in the macaque diet increased (16). During winter the macaques consumed leaves, stems, nuts, and bark, whereas in the summer they consumed principally mature leaves and stems in addition to corn, which was provisioned to habituate the monkeys to the presence of researchers (16). A previous study using DNA metabarcoding found that our population of common marmosets consumed more species of insects during the wetter period, but that the diversity of plants species consumed did not vary between wetter and drier periods (40). DNA metabarcoding cannot detect changes in plant part consumption, and the fungal communities of a single plant can vary by plant part (e.g. stem, seed, leaf, and root) (41). Caatinga marmosets are known to eat multiple parts from a single plant (32), so we cannot rule out the possibility that changes in plant part consumption between wetter and drier periods explain changes in their gut mycobiome composition between wetter and drier periods. To address this question, future investigators should sample the plant parts consumed and not consumed during the period of utilization and compare their fungal communities.

The five fungal ASVs that were differentially abundant between wetter and drier periods are known to be associated with plants as endophytes, epiphytes, and/or plant pathogens. Therefore, dietary differences between wetter and drier periods may explain the differentially abundant fungal ASVs. The more prevalent ASVs during the wetter period include two species of *Cladosporium*, which were more common in males, and are known plant pathogens (37). Another ASV that was associated with the wet period was *Candida parapsilosis* (phylum Ascomycota), which we classified as an animal pathogen, an endophyte, and a saprotroph. *Candida parapsilosis* is considered an important emerging fungal pathogen in humans, where it causes blood infections and is associated with hospital environments. It has been isolated from domestic animals and soil in addition to being considered a human skin commensal (42). We classified *Penicillium sumatraense* (phylum Ascomycota), associated with the wetter period, as a saprotroph, and it has also been described as an epiphyte and plant pathogen (43). We considered *Nothophoma*, which was more abundant during the drier period, to be a plant pathogen. We know that common marmosets at our site feed on multiple structures from the same species of plant (32), and evidence from other sites within the Caatinga biome shows variation in which plant parts are eaten throughout the year (44), which could explain the relationship between the marmoset mycobiome and climatic conditions. It is also possible that the fungal communities of the plants that the marmosets feed, travel, and rest on vary between wetter and drier periods. The variability of fungal taxa with likely dietary and/or environmental origins between wetter and drier periods suggests that the mycobiome is largely transient.

### Relationship between diet and mycobiome diversity

The positive correlation between within sample diversity metrics of the arthropod diet and the mycobiome indicates that a portion of the gut mycobiome may be transient. Arthropods harbor fungal pathogens (25) and consume plants that have their own fungal communities, which means that arthropods are a potential source of transient fungi. The lack of relationship between the within sample diversity of the plant portion of the marmoset diet and the overall mycobiome does not negate the possibility that much of the gut mycobiome is transient. Common marmosets may acquire some proportion of their mycobiota from the surface of trunks, branches, and leafy material as they forage, travel and rest in the arboreal canopy or from the ground during foraging and travel. In addition, the fact that we measured taxonomic diversity from plant DNA present in fecal samples, rather than the diversity of plant structures consumed (41), may have underrepresented the diversity of the common marmoset diet. Additionally, the significant correlation of the compositions of the mycobiome and the plant portion of the marmoset diet further supports a primarily transient mycobiome. The link between plant consumption and the gut mycobiome has been previously reported, with a 26-35% overlap between the mycobiota of the plant species consumed and fecal mycobiota in Tibetan macaques (18). We found that more than half of the FUNGuild-categorized ASVs in this study were associated with plants (plant pathogens, endophytes, epiphytes, bryophyte parasites, ectomycorrhizal fungi, or plant parasites). The correlation between the composition of the plant portion of the marmoset diet based on fecal DNA and their gut mycobiome may indicate that different species of plants contribute different fungal ASVs. In contrast, the lack of correlation between the number of plants and the number of fungal ASVs may indicate that the numbers of fungal ASVs associated with different plants varies. Correlations between the composition of the plant portion of the diet and prior work on the fungal communities of plants support the idea that the gut mycobiome of marmosets is largely transient.

### Is the gut mycobiome largely residential or primarily transient?

Common marmosets that consume a richer arthropod diet have richer transient fungal communities and richer and more even resident fungal communities. However, our results must be interpreted cautiously, as a low proportion of fungal taxa were classified to family or below (63%) or assigned to a functional guild (40%). In addition, since the marmosets consume both predatory and herbivorous arthropods, this could introduce transient fungal taxa associated with the plants that the arthropods consumed. Moreover, our resident fungal communities included animal pathogens, which could originate from arthropod prey. The within sample diversity of the plant portion of the diet did not correlate with the same metrics from the resident or transient portions of the mycobiome, which suggests that consuming a wider range of plants is not necessarily exposing the marmosets to a wider range of resident or transient fungi. This could suggest that the marmosets’ diet is both contributing to transient taxa and serving as a reservoir for the transmission of resident taxa.

We conclude that the common marmoset gut mycobiome may be largely transient. While we did find a significant relationship between a host factor (sex, males had greater within sample diversity than females) and within sample diversity metrics of the gut mycobiome (e.g. Shannon Diversity, Chao index, etc.) our results are compatible with the common marmoset gut mycobiome being more heavily shaped by external environmental factors (e.g. diet and climatic conditions) than by host factors. We found a significant relationship among the composition of the plant portion of the diet, wetter and drier periods of the year, and the composition of the gut fungal community, highlighting that these external factors were more closely related to the presence and absence of the mycobiota in the host gut. We also found that resident fungi in the marmoset gut represented a minority (8% of the fungi with ecological information) of the taxonomic composition of the gut mycobiome. The sex-based differences in the within sample diversity of the mycobiome may be best explained by different hormone levels between males and females, differences in substrate use, or differences in foraging habits associated with provisioning infants. Future work should investigate sex differences in behavior including substrate use, diet, and foraging strategies and directly sample the fungal communities of the common marmoset diet and the environment to more fully understand these findings.

## Materials and Methods

### Study Site

We studied eight groups of common marmosets at the Baracuhy Biological Field Station (7°31’42’’S, 36°17’50’’W) in the state of Paraíba, Brazil. This site has a hot semi-arid climate with thorn scrub vegetation and distinct wet and dry periods. Average rainfall is ∼337 mm per year with records showing some years with almost no rainfall (29,45). For seven months of the year daytime temperatures may surpass 35°C, with the average maximum temperature being 29.1°C during the wetter period and 31.3°C in the drier period (29,32). The marmosets used in this study were part of an experimental field study of decision-making, food choice, and access to resources. To accomplish this, we placed artificial feeding stations within each group’s home range that contained bananas and meal worms. We baited the feeding stations three times daily and varied in the amount of food provided (See 27 and 29 for more detailed information on provisioning). Once the marmosets left the feeding station they fed on their natural diet.

### Sample Collection

We collected common marmoset fecal samples (N = 67) across the wetter (February and March) and drier (July and August) periods of 2015 and 2016 as described elsewhere (40). We trapped the marmosets, determined their sex, estimated their approximate age, marked them with individual color-coded beaded collars, and released them unharmed (40). We collected feces from pans set beneath traps housing individual marmosets and stored in tubes of either RNAlater or 95% ethanol (40).

### DNA Extraction

DNA extraction followed existing protocols (40,46). Briefly, we extracted DNA from 67 fecal samples using the Qiagen PowerSoil Kit with two protocol modifications: we incubated samples for 15 minutes at 65°C after adding solution C1 and before bead beating. We incubated samples at room temperature for 5 minutes with solution C6 having been pre-warmed to 65°C prior to centrifugation. Extraction negatives, which used molecular grade water in place of feces, were included to control for contamination during the extraction procedure (40).

### Mycobiome sequencing and analyses

We amplified the ITS3 and ITS4 regions of the ITS rDNA region for our samples with the ITS3 (5’-GCATCGATGAAGAACGCAGC) - ITS4 (5’-TCCTCCGCTTATTGATATGC) primer set on the Fluidigm Access Array system as previously described (40,47). We used the cutadapt function to trim our sequences due to the variable length of the ITS amplicon. We used the DADA2 workflow in QIIME2 (v. 2023.9.1) to quality filter and denoise the rDNA ITS sequences into amplicon sequence variants (ASVs) (48). We created a naïve Bayesian classifier using the UNITE database (version 9.999 downloaded on Oct. 27, 2022) in QIIME and used it to assign taxonomy to the ASVs. We rarefied samples to 5,000 reads based on rarefaction plots (Figure S10). We removed seven samples (Table S33) that had fewer than 5,000 reads from further within sample diversity and composition analyses. We subsampled the 60 retained samples 1,000 times to 5,000 reads, and we calculated the Shannon diversity, Simpson diversity, Chao1 diversity, ASV richness, and Pielou’s evenness metrics for each subsample. We recorded the average value of each metric as the rarefied value of that metric for that sample. We used the *vegan* package (v. 2.7-1) in R version 4.4.0 (49,50) to calculate the Shannon and Simpson diversity indices (1 – D). We calculated the Chao1 diversity using the *fossil* package (v. 0.4.0) (51). We calculated the richness by counting the observed features, and calculated Pielou’s evenness (52). We omitted two samples that had a disproportionate impact on the non-metric multidimensional scaling plots (MDS1 values <-3 and MDS2 values <-15) from subsequent composition analyses (resulting n = 50; Tables S34-35) to avoid these samples skewing results. We used a single 5,000-read subsample of each sample to calculate the mycobiome composition as Bray-Curtis and Jaccard distances with the *vegan* package (v 2.7-1) (49).

### FUNGuild Ecological Role Assignment and analyses

We used FUNGuild (v. 1.2) (26), a tool that can sort fungal ASVs into guilds based on their ecological roles, to sort ASVs with sufficient taxonomic information into guilds. We considered some guilds (i.e. Plant Pathogen, Endophyte, Epiphyte, Bryophyte Parasite, Ectomycorrhizal, Plant Parasite) to be likely transient taxa associated with the marmoset diet and other guilds (i.e. Animal Pathogen, Endosymbiont, Animal Endosymbiont) to be likely resident taxa. Fungal taxa with guilds in both categories was considered a transient taxon (Figure S11) due to relationships between plants and fungi being more common than animal/fungus relationships (53) and many fungi being opportunistically pathogenic for humans (54). We calculated the within sample diversity metrics of these two categories as described above for the overall mycobiome.

### Diet metabarcoding and analysis

We characterized the marmoset diet using metabarcoding techniques as described in Finnegan et al. (2024), and using different markers for different portions of the diet. We used the trnLg-trnLh primer pair to amplify the *trnL* gene to characterize the plant portion of the diet, and the ArtF11-ArtR17 primer pair, which amplifies the *COI* gene, for the arthropod portion (55). We amplified samples with a Fluidigm Access Array system, performed PCR with standard protocols (47), and sequenced samples on the Illumina NovaSeq platform with a 2×250 bp SP flowcell at the Roy J. Carver Biotechnology Center at the University of Illinois Urbana-Champaign to generate reads (40). We used Obitools3 (v. 3.0.0b33) to pair, quality filter, and assign taxonomy to the reads (56). We constructed plant and arthropod referenced databases from the EMBL nucleotide sequence database (downloaded August 17, 2020) with ecoPCR as implemented in Obitools3 (https://github.com/emallott/Caatinga_marmosets) (40,56). We calculated Shannon diversity, Simpson diversity, Chao1 diversity, richness (i.e. observed features), and Pielou’s evenness to assess within sample diversity of the plant and arthropod portions of the diet. We calculated Bray-Curtis and Jaccard distances using the *vegan* package (v 2.7-1) (49) to assess sample composition.

### Statistical analyses

We carried out data analyses in R (v. 4.4.0) (50) and RStudio (v. 2025.5.0.496) (57). We used linear models to determine if host sex and/or sample collection period (wet or dry) correlated with the within sample diversity metrics that were calculated for the marmoset diet and mycobiome. We omitted samples that did not have an assigned sex from these analyses (resulting in n = 52; Table S36). We used linear mixed-effect models, as implemented in the *lme4* package in R (v 1.1-37) (58) to assess the effects of host sex and age, sample collection period, sample preservative (ethanol or RNAlater), and sample collection time (a combination of period and year). We included social group as a random effect. When the mixed-effect model was singular, we used a fixed-effect modeling approach that included social group as an additional fixed effect (Table S37). We determined significance of the parameters of interest in mixed-effect models using likelihood ratio tests where a full model was compared to a null model missing the parameter of interest. We used the *stats* package (v. 4.4.0) (50) to run analysis of variance (ANOVA) to compare the full and null models. We determined the significance of the difference between levels of a parameter (e.g. male and female for sex) by running the Wald’s test in the *car* package (v. 3.1-3) (59).

We used permutational multivariate analysis of variance (PERMANOVA), as implemented in the *vegan* package (v. 2.7-1) (49), to determine whether host sex and/or sample collection period varied with the mycobiome composition metrics Bray-Curtis and Jaccard distances for the diet and mycobiome. We visualized these results using non-metric multidimensional scaling (NMDS). We calculated the variance of the sample categories using multivariate homogeneity of groups dispersions, both using the *vegan* package (49). We compared the variances of the sample categories with analysis of variance (ANOVA) as implemented in the *stats* package (50).

We used a full set of samples with sufficient read counts (n = 60) to test the associations between diet and mycobiome. We tested interactions between the within sample diversity metrics of the mycobiome and the arthropod and plant portions of the diet using Spearman rank correlation tests as implemented in *stats* package (50). We tested interactions between these two portions of the diet and the overall mycobiome, the transient mycobiome, and the resident mycobiome.

We tested interactions between diet and mycobiome composition using Mantel tests as implemented in the *ecodist* package (v. 2.1.3) (60). We used analysis of compositions of microbiomes with bias correction (ANCOM-BC), as implemented in the *ANCOMBC* package (v. 2.6.0) (61,62), to identify ASVs whose abundance differs between sample collection periods and between sexes. We did not drop samples with fewer than 5,000 reads, as ANCOM-BC accounts for differential library sizes, but we did drop samples without sex information (n = 57). We used a Benjamini-Hochberg procedure to reduce the false discovery rate. We tested interactions between specific dietary and fungal ASVs using compositionality corrected renormalization and permutation, which uses a Benjamin-Hochberg-Yekutieli procedure to control false discovery rate, as implemented in the *ccrepe* package (v. 1.40.0) (63) following the example of Sharma et al. (2022). We visualized these results with Cytoscape (v 3.10.1) (64).

## Data Availability

Sequences are available in the Short Read Archive (http://www.ncbi.nlm.nih.gov/sra) under BioProject PRJNA1021612. All metadata used in the analyses and code for all analyses can be found on GitHub (https://github.com/Mallott-Lab/CjacchusITS).

## Acknowledgements

The authors are grateful to Dr Geraldo Baracuhy for allowing us to conduct this study at the Baracuhy Biological Field Station. The field component and sample collection of this study were supported by the Brazilian Higher Education Authority/CAPES (grant/award PVE # 88881.064998/2014-01). PAG acknowledges the support provided by Chrissie, Sara, Jenni, Dax, and Saffron during the writing of this manuscript. JCBM acknowledges the Brazilian National Council for Scientific and Technological Development/CNPq for research fellowships (PQ #303306/2013-0 and 311731/2023-6). KRA is supported as a fellow of CIFAR’s ‘Humans and the Microbiome’ group. NS acknowledges the Conselho Nacional de Desenvolvimento Científico e Tecnológico, grant/award number: APQ 403126/2016-9.

B.J.G. and E.K.M. conceived and designed the study. P.A.G., A.C.M., M.F.D.F., F.A., A.S., and N.S. conducted field data collection. P.M.F. and E.K.M. conducted the lab work. B.J.G. and E.K.M. analyzed the sequencing data and carried out the statistical analysis. B.J.G. drafted the manuscript. All authors read, edited, and approved the final manuscript.

