## Supplemental Information for "Diet, Climatic Conditions, and Sex Affect the Mycobiome of Wild Common Marmosets (*Callithrix jacchus*)"

Figure S1. Boxplots comparing the Chao1 diversity of samples by sex (A) and sampling period (B). Wald's tests showed that males had significantly higher Chao1 diversity values (p = 0.009005; Table S3). There was no significant difference between the dry and wet sample collection periods (p = 0.232757; Table S3). Boxplots comparing the Shannon diversity of the samples by sex (C) and sampling period (D). Wald's tests showed that males had significantly higher Shannon diversity values (p = 0.0005105; Table S1). There was no significant difference between the dry and wet sample collection periods (p = 0.1728822; Table S1)........ 5

Figure S5. Non-metric multidimensional scaling (NMDS) plot showing the grouping of samples, which are colored by sample collection period. Each point represents a single sample. Ellipses represent the standard deviation of the points within a group (Oksanen et al., 2022). The plots were generated using the Jaccard distances of the mycobiome. The PERMANOVA shows that the effect of sample collection period (pseudo-F(1) = 1.1776, p = 0.006; Table S7) was statistically significant. .......................................10

Figure S6. Non-metric multidimensional scaling (NMDS) plot showing the grouping of samples, colors and shapes indicate social group. Each point represents a single sample. Ellipses represent the standard deviation of the points within a group (Oksanen et al., 2022). The plots were generated using the Jaccard distances of the mycobiome. The PERMANOVA shows that the effect of social group (pseudo-F(7) = 1.0548, p = 0.013; Table S7) was statistically significant. .......................................11

Table S16. The results of a fixed-effects model that tested which parameter levels impacted the Simpson diversity of the likely resident fungi. No parameters were significant, and the overall model was not significant (p = 0.6564). 16

Table S17. The results of a fixed-effects model that tested which parameter levels impacted the Pielou's evenness values of the likely resident fungi. No parameters were significant, and the overall model was not significant (p = 0.5152). 16

Table S18. The results of the ANCOM-BC, indicating which ASVs were differentially abundant between the sample collection periods. The drier period is the reference, with these values being associated with the wetter period. 17

Table S19. The results of several Spearman correlation tests comparing the within sample diversity metrics of the plant portion of the marmoset diet to the corresponding metrics associated with the overall mycobiome, the likely transient fungi, and the likely resident fungi. None of the correlations were statistically significant. 17

Table S20. The results of several Spearman correlation tests comparing the within sample diversity metrics of the arthropod portion of the marmoset diet to the corresponding metrics associated with the overall mycobiome, the likely transient fungi, and the likely resident fungi. Significant correlations are bolded. 18

Table S21. The results of a CCREPE analysis investigating interactions between 12 fungal ASVs and 44 arthropod ASVs. Each row represents a pairwise interaction, with columns showing the fungal ASV, arthropod ASV, p-value, q-value, sim score, and z stat. Table is included as a separate file titled SupplementalTable21.xlsx 18

Figure S8. Network of interactions between fungal ASVs (red) and arthropod ASVs (purple). No interaction was significant (q-value < 0.05 and absolute sim score > 0.6; Table S21). Fungal ASVs that were identified by ANCOM-BC are labeled. 19

Table S22. The results of a CCREPE analysis investigating interactions between 12 fungal ASVs and 266 plant ASVs. Each row represents a pairwise interaction, with columns showing the fungal ASV, arthropod ASV, p-value, q-value, sim score, and z stat. The significant relationship was bolded and moved to the top of the table. Table is included as a separate file titled SupplementalTable22.xlsx 19

Figure S9. Network of interactions between fungal ASVs (red) and plant ASVs (gray). Only ASVs involved in interactions with q-values below 1 were mapped. Significant interactions are indicated by thick bolded edges (Table S22). ASVs that were identified as significant by ANCOM-BC and/or CCREPE are labeled. 20

Table S23. Results of likelihood ratio tests comparing full mixed-effects models to null models missing the sex parameter to determine whether it significantly relates to within sample diversity metrics of arthropod portion of the arthropod diet. None of the models were significant. 20

Table S24. The results of a fixed-effects model that tested which parameter levels related to the Pielou's evenness of the arthropod portion of the marmoset diet. None of the parameters were significant. 21

Table S25. The results of a fixed-effects model that tested which parameter levels related to the Shannon diversity of the plant portion of the marmoset diet. None of the parameters were significant. 21

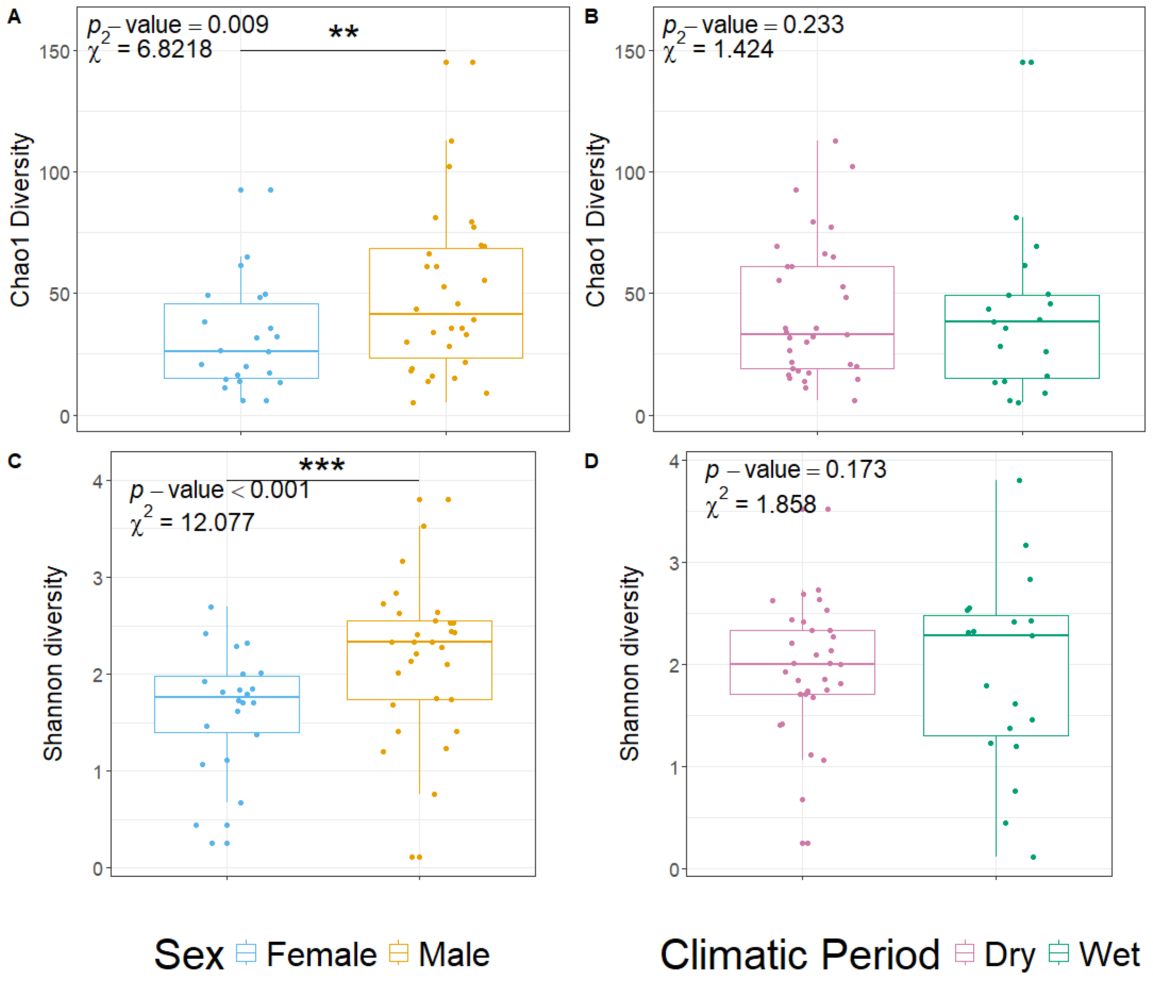

**Figure S1.** Boxplots comparing the Chao1 diversity of samples by sex (A) and sampling period (B). Wald's tests showed that males had significantly higher Chao1 diversity values (p = 0.009005; Table S3). There was no significant difference between the dry and wet sample collection periods (p = 0.232757; Table S3). Boxplots comparing the Shannon diversity of the samples by sex (C) and sampling period (D). Wald's tests showed that males had significantly higher Shannon diversity values (p = 0.0005105; Table S1). There was no significant difference between the dry and wet sample collection periods (p = 0.1728822; Table S1).

**Table S1.** The results of a Wald's test of a mixed-effects model that tested which parameters significantly impacted the Shannon diversity value. Significant parameters are bolded.

| Parameter | Chi Squared | Degrees of Freedom | p-value |
|---|---|---|---|
| Sampling Period | 1.8578 | 1 | 0.1728822 |
| **Sex** | **12.0771** | **1** | **0.0005105** |
| Preservative | 0.3668 | 1 | 0.5447633 |
| Age | 7.4004 | 3 | 0.0601739 |
| Time | 0.5773 | 2 | 0.7492596 |

**Table S2.** The results of a Wald's test of a mixed-effects model that tested which parameters significantly impacted the richness value. Significant parameters are bolded.

| Parameter | Chi Squared | Degrees of Freedom | p-value |
|---|---|---|---|
| Sampling Period | 1.1075 | 1 | 0.292616 |
| **Sex** | **6.8275** | **1** | **0.008976** |
| Preservative | 0.0066 | 1 | 0.935399 |
| Age | 5.0935 | 3 | 0.16508 |
| Time | 1.2393 | 2 | 0.538128 |

**Table S3.** The results of a Wald's test of a mixed-effects model that tested which parameters significantly impacted the Chao1 value. Significant parameters are bolded.

| Parameter | Chi Squared | Degrees of Freedom | p-value |
|---|---|---|---|
| Sampling Period | 1.4239 | 1 | 0.232757 |
| **Sex** | **6.8218** | **1** | **0.009005** |
| Preservative | 0.0111 | 1 | 0.916246 |
| Age | 5.2147 | 3 | 0.150154 |
| Time | 1.5133 | 2 | 0.469247 |

**Table S4.** The results of a fixed-effects model that tested which parameter levels impacted the Simpson diversity value. Significant parameter levels are bolded.

| Parameter-Level | Estimate | Standard Error | t value | p-value |
|---|---|---|---|---|
| **Period-Wet** | **-0.178905** | **0.083454** | **-2.144** | **0.03851** |
| **Sex-Male** | **0.167276** | **0.059453** | **2.814** | **0.00772** |
| Group-Coqueiro | -0.135629 | 0.133467 | -1.016 | 0.31596 |
| Group-Cow | -0.152864 | 0.178027 | -0.859 | 0.39591 |
| Group-F Group | -0.019503 | 0.162084 | -0.12 | 0.90486 |
| Group-House | -0.043259 | 0.116274 | 0.372 | 0.71193 |
| Group-Key | 0.183898 | 0.144044 | 1.277 | 0.20946 |
| Group-Princess | -0.08966 | 0.115184 | -0.778 | 0.44115 |

| | | | | |
|---|---|---|---|---|
| Group-Road | 0.028378 | 0.14775 | 0.192 | 0.84871 |
| Preservative-RNAlater | 0.006245 | 0.075572 | 0.083 | 0.93457 |
| Age-Infant | -0.129144 | 0.127488 | -1.013 | 0.31747 |
| Age-Juvenile | -0.11297 | 0.086303 | -1.309 | 0.1984 |
| **Age-Subadult** | **0.30975** | **0.140808** | **2.2** | **0.03398** |

**Table S5.** The results of a fixed-effects model that tested which parameter levels impacted the Pielou’s evenness value. Significant parameter levels are bolded.

| **Parameter-Level** | **Estimate** | **Standard Error** | **t value** | **p-value** |
|---|---|---|---|---|
| Period-Wet | -0.10772 | 0.06363 | -1.693 | 0.09862 |
| **Sex-Male** | **0.12293** | **0.04533** | **2.712** | **0.00998** |
| Group-Coqueiro | -0.06966 | 0.10176 | -0.685 | 0.49777 |
| Group-Cow | -0.12082 | 0.13573 | -0.89 | 0.37899 |
| Group-F Group | -0.01176 | 0.12357 | -0.95 | 0.92467 |
| Group-House | -0.02575 | 0.08865 | -0.291 | 0.77301 |
| Group-Key | 0.16771 | 0.10982 | 1.527 | 0.13501 |
| Group-Princess | -0.02164 | 0.08782 | -0.246 | 0.80665 |
| Group-Road | 0.03414 | 0.11265 | 0.303 | 0.7635 |
| Preservative-RNAlater | -0.01759 | 0.05762 | -0.305 | 0.76182 |
| Age-Infant | -0.13041 | 0.0972 | -1.342 | 0.18766 |
| Age-Juvenile | -0.06948 | 0.0658 | -1.056 | 0.29767 |
| **Age-Subadult** | **0.23998** | **0.10735** | **2.235** | **0.03134** |

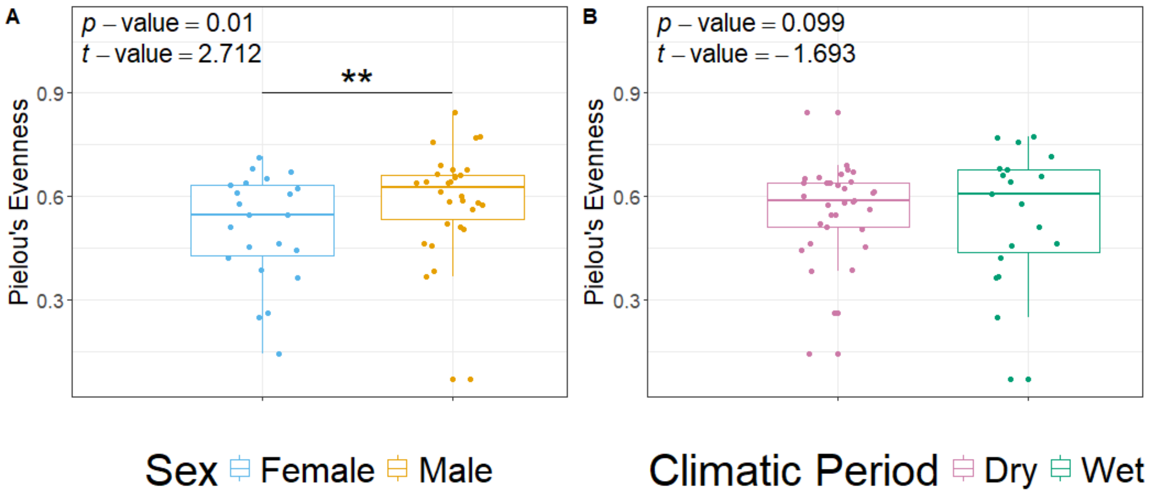

**Figure S2.** Boxplots comparing the Pielou's evenness of samples by sex (A) and sampling period (B). Fixed-effects models show that Pielou's evenness varied by sex (p = 0.0098, t-value = 2.712, Table S5) but not sampling period (p = 0.09862, t-value = -1.693, Table S5).

**Table S6.** The results of a PERMANOVA using marginal sums of squares looking at the effects of various parameters on the Bray-Curtis distances between the mycobiome compositions of common marmoset samples. Significant parameters are bolded (Figure 2E-F; Figures S3-4). This analysis is informed by 50 samples. The variance of the samples did not differ between the sexes (p = 0.8101) or sampling periods (p = 0.831) for the Bray-Curtis distance.

| Parameter | Degrees of Freedom | Sum of Squares | $R^2$ | Pseudo-F | p-value |
|---|---|---|---|---|---|
| Age | 3 | 1.0804 | 0.05131 | 0.9087 | 0.711 |
| Sex | 1 | 0.4123 | 0.01958 | 1.0404 | 0.358 |
| **Period** | **1** | **0.5844** | **0.02776** | **1.4746** | **0.048** |
| **Group** | **7** | **3.5418** | **0.16822** | **1.2767** | **0.014** |
| **Preservative** | **1** | **0.8358** | **0.0397** | **2.1089** | **0.002** |
| Residuals | 36 | 14.2668 | 0.67763 | | |

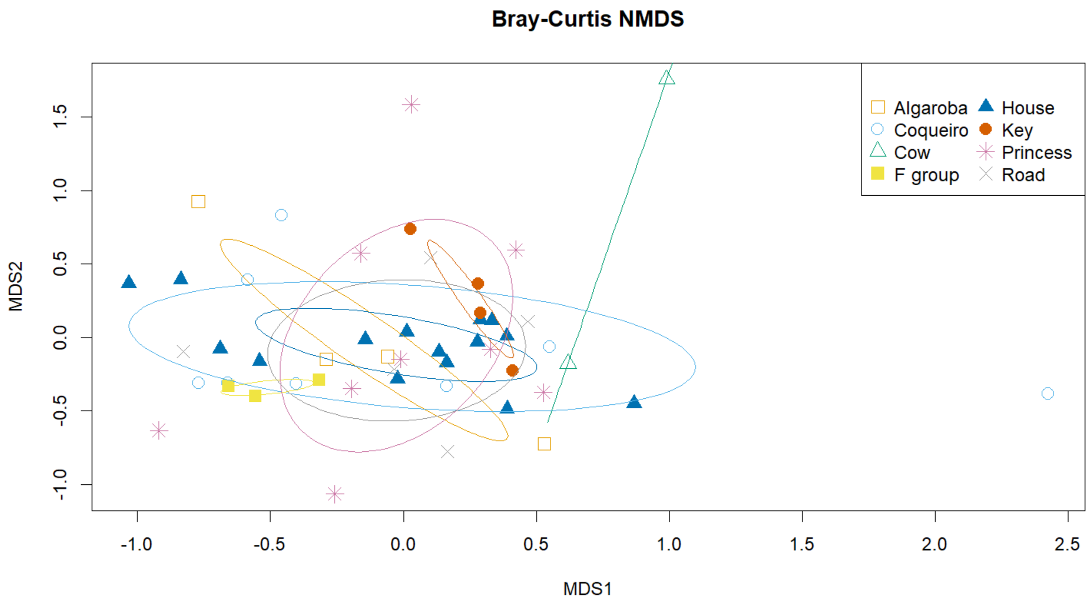

**Figure S3.** Non-metric multidimensional scaling (NMDS) plot showing the grouping of samples, colors and shapes indicate social group. Each point represents a single sample. Ellipses represent the standard deviation of the points within a group (Oksanen et al., 2022). The plots were generated using the Bray-Curtis distances of the mycobiome. The PERMANOVA shows that the effect of social group (pseudo-F(7) = 1.2767, p = 0.014; Table S6) was statistically significant.

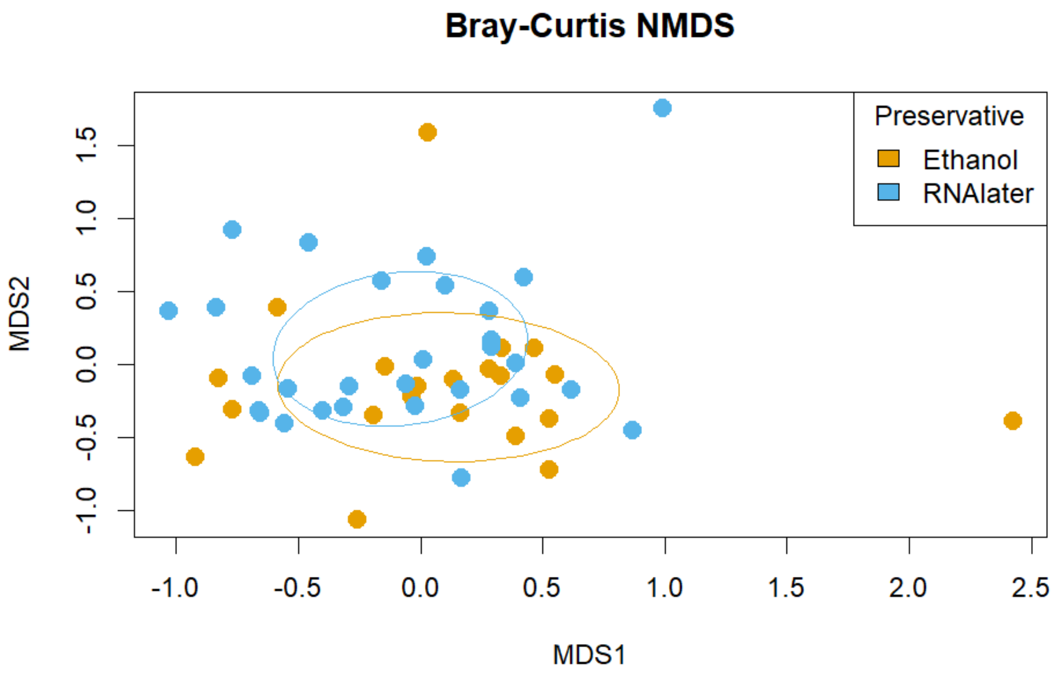

**Figure S4.** Non-metric multidimensional scaling (NMDS) plot showing the grouping of samples, which are colored by preservative. Each point represents a single sample. Ellipses represent the standard deviation of the points within a group (Oksanen et al., 2022). The plots were generated using the Bray-Curtis distances of the mycobiome. The PERMANOVA shows that the effect of social group (pseudo-F(1) = 2.1089, p = 0.002; Table S6) was statistically significant.

**Table S7.** The results of a PERMANOVA using marginal sums of squares looking at the effects of various parameters on the Jaccard distances between the mycobiome compositions of common marmoset samples. Significant parameters are bolded (Figures S5-7). This analysis is informed by 50 samples. The variance of the samples did not differ between the sexes (p = 0.1619) or sampling periods (p = 0.083) for the Jacard distance.

| Parameter | Degrees of Freedom | Sum of Squares | $R^2$ | Pseudo-F | p-value |
|---|---|---|---|---|---|
| Age | 3 | 1.4524 | 0.06216 | 1.0333 | 0.151 |
| Sex | 1 | 0.4936 | 0.02112 | 1.0535 | 0.141 |
| **Period** | **1** | **0.5517** | **0.02361** | **1.1776** | **0.006** |
| **Group** | **7** | **3.4594** | **0.14805** | **1.0548** | **0.013** |
| **Preservative** | **1** | **0.5525** | **0.02365** | **1.1793** | **0.008** |
| Residuals | 36 | 16.8662 | 0.72181 | | |

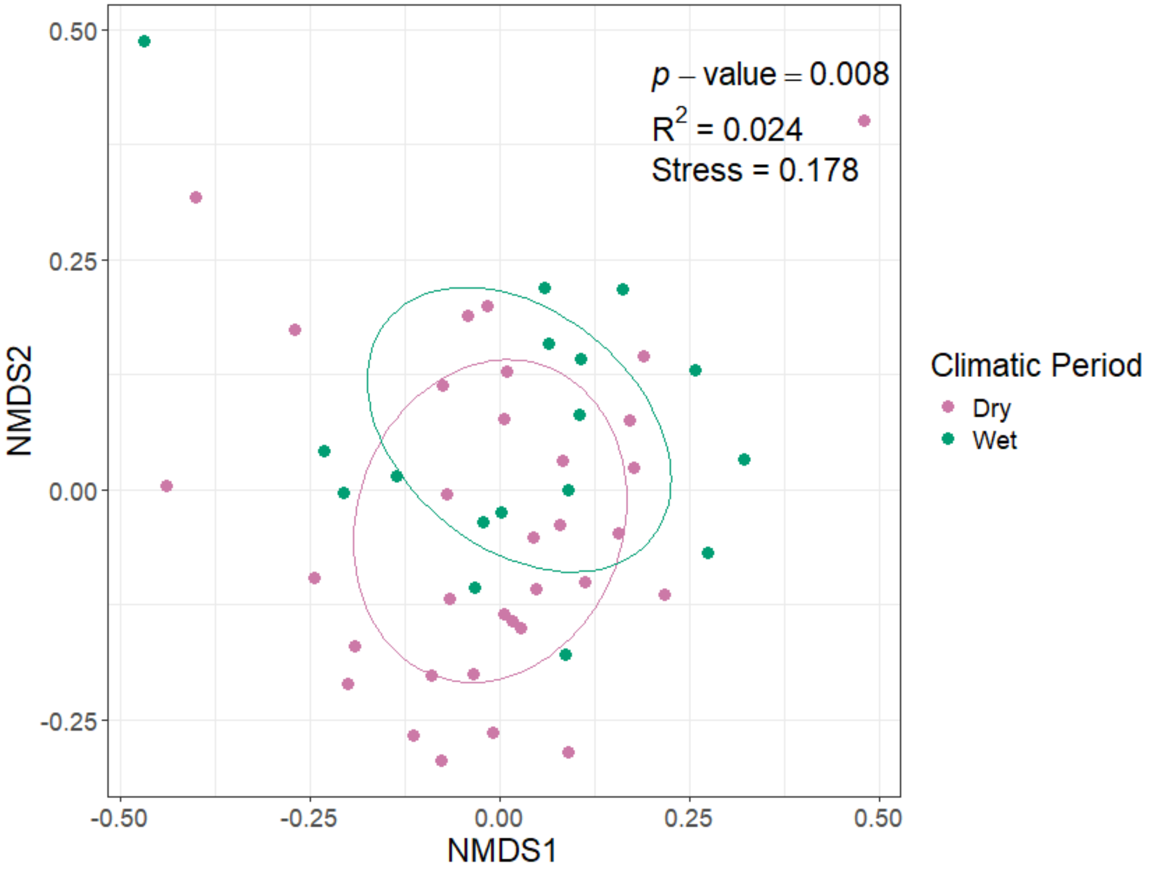

**Figure S5.** Non-metric multidimensional scaling (NMDS) plot showing the grouping of samples, which are colored by sample collection period. Each point represents a single sample. Ellipses represent the standard deviation of the points within a group (Oksanen et al., 2022). The plots were generated using the Jaccard distances of the mycobiome. The PERMANOVA shows that the effect of sample collection period (pseudo-F(1) = 1.1776, p = 0.006; Table S7) was statistically significant.

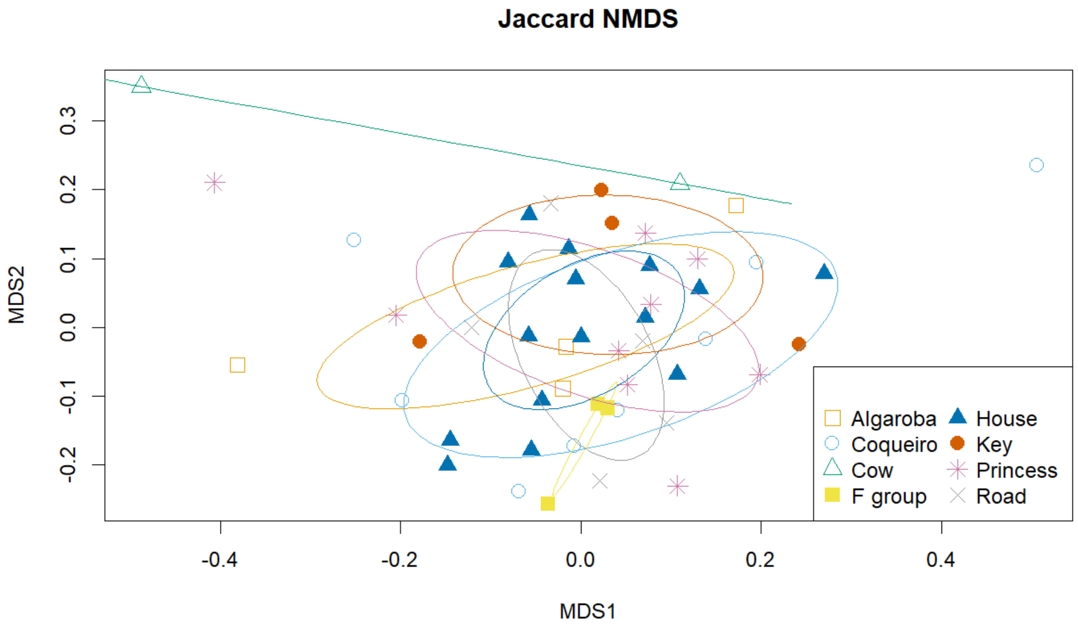

**Figure S6.** Non-metric multidimensional scaling (NMDS) plot showing the grouping of samples, colors and shapes indicate social group. Each point represents a single sample. Ellipses represent the standard deviation of the points within a group (Oksanen et al., 2022). The plots were generated using the Jaccard distances of the mycobiome. The PERMANOVA shows that the effect of social group (pseudo-F(7) = 1.0548, p = 0.013; Table S7) was statistically significant.

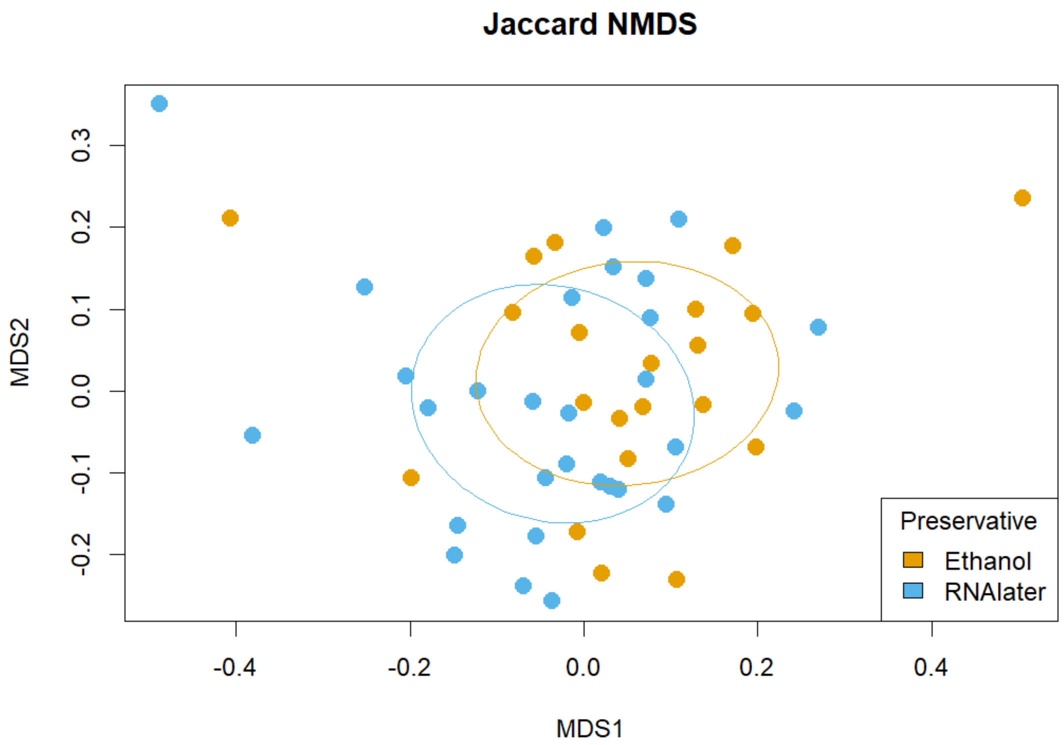

**Figure S7.** Non-metric multidimensional scaling (NMDS) plot showing the grouping of samples, which are colored by preservative. Each point represents a single sample. Ellipses represent the standard deviation of the points within a group (Oksanen et al., 2022). The plots were generated using the Jaccard distances of the mycobiome. The PERMANOVA shows that the effect of social group (pseudo-F(1) = 1.1793, p = 0.008; Table S7) was statistically significant.

**Table S8.** The results of the ANCOM-BC, indicating which ASVs were differentially abundant between the sexes. The female sex is the reference, with these values being associated with the male sex.

| ASV | Taxon | LFC | Standard Error | p-value |
|---|---|---|---|---|
| ff0d19a450862c945b591b0c7f6d5181 | Fungus 1 | -1.330 | 0.346 | 0.005 |
| 09747f28421a8d60ef07b64fdd7263b0 | *Nothophoma* sp. | -2.413 | 0.472 | 0.005 |
| e462722cb5bc4b112adfc8babf18e384 | *Cladosporium* 1 | 2.171 | 0.255 | 0.003 |
| 9839ecfe0e871175e6b0acd6bcc64204 | *Leptosillia* sp. | -2.858 | 0.228 | 0.005 |
| 18aeb211ab36e34ea362709b418599cb | *Cladosporium* 2 | 1.732 | 0.162 | 0.007 |
| 9d8a544f6e722db95e04df5a787029a2 | Fungus 2 | -2.040 | 0.186 | 0.005 |
| 66ba111b8604449b0bd5840acc15b401 | Fungus 3 | -1.062 | 0.173 | 0.012 |

**Table S9.** Results of likelihood ratio tests comparing full mixed-effects models to null models missing the parameter of interest, sex or sampling period, to determine whether they have significant impacts on within sample diversity metrics of the likely transient fungi. Significant models are bolded.

| Diversity Metric | Model | logLik | deviance | Chisq | Df | Pr(>Chisq) |
|---|---|---|---|---|---|---|
| ASV Richness | Full Model | -163.13 | 326.26 | 0 | 1 | 0 |
| | Null Climatic Period Model | -163.13 | 326.26 | | | |
| | **Full Model** | **-163.13** | **326.26** | **6.462** | **1** | **0.01102** |
| | Null Sex Model | -166.36 | 332.72 | | | |
| Chao1 | Full Model | -170.45 | 340.9 | 0 | 0 | |
| | Null Climatic Period Model | -170.45 | 340.9 | | | |
| | **Full Model** | **-170.45** | **340.9** | **6.1799** | **1** | **0.01292** |
| | Null Sex Model | -173.54 | 347.08 | | | |

**Table S10.** The results of a fixed-effects model that tested which parameter levels impacted the Shannon diversity value of the likely transient fungi. Significant parameter levels are bolded. The overall model was not significant (p = 0.3412).

| Parameter-Level | Estimate | Standard Error | t value | p-value |
|---|---|---|---|---|
| Period-Wet | -0.3437 | 0.3049 | -1.127 | 0.2668 |
| Sex-Male | 0.3466 | 0.2172 | 1.596 | 0.1189 |
| Group-Coqueiro | -0.2687 | 0.4877 | -0.551 | 0.5849 |
| Group-Cow | -0.52688 | 0.6505 | -0.81 | 0.4231 |
| Group-F Group | 0.1179 | 0.5922 | 0.199+ | 0.8433 |
| Group-House | -0.3558 | 0.4249 | -0.837 | 0.4076 |
| Group-Key | 0.2022 | 0.5263 | 0.384 | 0.703 |
| Group-Princess | -0.6664 | 0.4209 | -1.583 | 0.1217 |
| Group-Road | 0.1164 | 0.5399 | 0.216 | 0.8305 |
| Preservative-RNAlater | 0.2855 | 0.2761 | 1.034 | 0.3078 |
| Age-Infant | -0.1027 | 0.4658 | -0.22 | 0.8267 |
| Age-Juvenile | -0.6104 | 0.3153 | -1.936 | 0.0604 |
| Age-Subadult | 0.2213 | 0.5145 | 0.43 | 0.6695 |

**Table S11.** The results of a fixed-effects model that tested which parameter levels impacted the Simpson diversity value of the likely transient fungi. Significant parameter levels are bolded. The overall model was not significant (p = 0.9607).

| Parameter-Level | Estimate | Standard Error | t value | p-value |
|---|---|---|---|---|
| Period-Wet | -0.058111 | 0.141258 | -0.411 | 0.6831 |

| | | | | |
|---|---|---|---|---|
| Sex-Male | 0.121445 | 0.100633 | 1.207 | 0.235 |
| Group-Coqueiro | 0.041743 | 0.225912 | 0.185 | 0.8544 |
| Group-Cow | 0.236678 | 0.301337 | 0.785 | 0.4371 |
| Group-F Group | 0.152868 | 0.274351 | 0.557 | 0.5807 |
| Group-House | 0.032388 | 0.196811 | 0.165 | 0.8702 |
| Group-Key | 0.05024 | 0.243816 | 0.206 | 0.8378 |
| Group-Princess | -0.10392 | 0.194966 | -0.533 | 0.5971 |
| Group-Road | 0.003135 | 0.250088 | 0.013 | 0.9901 |
| Preservative-RNAlater | 0.00686 | 0.127917 | 0.054 | 0.9575 |
| Age-Infant | -0.202962 | 0.215792 | -0.941 | 0.3529 |
| Age-Juvenile | -0.173767 | 0.146081 | -1.19 | 0.2416 |
| Age-Subadult | 0.169932 | 0.238339 | 0.713 | 0.4802 |

**Table S12.** The results of a fixed-effects model that tested which parameter levels impacted the Pielou's evenness value of the likely transient fungi. Significant parameter levels are bolded. The overall model was not significant (p = 0.5116).

| **Parameter-Level** | **Estimate** | **Standard Error** | **t value** | **p-value** |
|---|---|---|---|---|
| Period-Wet | 0.1032 | 0.14647 | 0.705 | 0.487 |
| Sex-Male | 0.06477 | 0.10322 | 0.628 | 0.536 |
| Group-Coqueiro | 0.05487 | 0.218411 | 0.251 | 0.804 |
| Group-Cow | -0.35389 | 0.25657 | -1.379 | 0.179 |
| Group-F Group | 0.25144 | 0.23863 | 1.054 | 0.301 |
| Group-House | 0.0658 | 0.18586 | 0.354 | 0.726 |
| Group-Key | 0.10492 | 0.22651 | 0.463 | 0.647 |
| Group-Princess | 0.21797 | 0.21786 | 1.001 | 0.326 |
| Group-Road | 0.1212 | 0.2224 | 0.545 | 0.59 |
| Preservative-RNAlater | 0.19503 | 0.11909 | 1.638 | 0.113 |
| Age-Infant | 0.08599 | 0.22718 | 0.379 | 0.708 |
| Age-Juvenile | -0.17039 | 0.13115 | -1.299 | 0.205 |
| Age-Subadult | -0.05838 | 0.31517 | -0.185 | 0.854 |

**Table S13.** The results of a fixed-effects model that tested which parameter levels impacted the Shannon diversity value of the likely resident fungi. Significant parameter levels are bolded. The overall model was not significant (p = 0.4309).

| **Parameter-Level** | **Estimate** | **Standard Error** | **t value** | **p-value** |
|---|---|---|---|---|
| Period-Wet | 0.02618 | 0.15785 | 0.166 | 0.8691 |
| **Sex-Male** | **0.23287** | **0.11246** | **2.071** | **0.0452** |
| Group-Coqueiro | 0.03801 | 0.25245 | 0.151 | 0.8811 |

| | | | | |
|---|---|---|---|---|
| Group-Cow | -0.29916 | 0.33674 | -0.888 | 0.3799 |
| Group-F Group | -0.22992 | 0.30658 | -0.75 | 0.4579 |
| Group-House | -0.05248 | 0.21993 | -0.239 | 0.8127 |
| Group-Key | 0.01829 | 0.27246 | 0.067 | 0.94668 |
| Group-Princess | -0.24463 | 0.21787 | -1.123 | 0.2686 |
| Group-Road | -0.31288 | 0.27947 | -1.12 | 0.2699 |
| Preservative-RNAlater | -0.04598 | 0.14295 | -0.322 | 0.7495 |
| Age-Infant | 0.09621 | 0.24114 | -0.399 | 0.6922 |
| Age-Juvenile | -0.04808 | 0.16324 | -0.295 | 0.77 |
| Age-Subadult | -0.04088 | 0.26634 | -0.153 | 0.8788 |

**Table S14.** The results of a fixed-effects model that tested which parameter levels impacted the ASV richness of the likely resident fungi. No parameters were significant, and the overall model was not significant (p = 0.882).

| Parameter-Level | Estimate | Standard Error | t value | p-value |
|---|---|---|---|---|
| Period-Wet | -0.01877 | 0.6247 | -0.03 | 0.976 |
| Sex-Male | 0.62987 | 0.44504 | 1.415 | 0.165 |
| Group-Coqueiro | -0.06848 | 0.99908 | -0.069 | 0.946 |
| Group-Cow | -0.92434 | 1.33264 | -0.694 | 0.492 |
| Group-F Group | -1.00399 | 1.2133 | -0.827 | 0.413 |
| Group-House | -0.18914 | 0.87038 | -0.217 | 0.829 |
| Group-Key | 0.45509 | 1.07826 | 0.422 | 0.675 |
| Group-Princess | -0.60872 | 0.86222 | -0.706 | 0.485 |
| Group-Road | -0.54074 | 1.106 | -0.489 | 0.628 |
| Preservative-RNAlater | -0.23134 | 0.56571 | -0.409 | 0.685 |
| Age-Infant | 0.27945 | 0.95433 | 0.293 | 0.771 |
| Age-Juvenile | -0.32943 | 0.64603 | -0.51 | 0.613 |
| Age-Subadult | -0.0403 | 1.05404 | -0.038 | 0.97 |

**Table S15.** The results of a fixed-effects model that tested which parameter levels impacted the Chao1 diversity of the likely resident fungi. No parameters were significant, and the overall model was not significant (p = 0.8856).

| Parameter-Level | Estimate | Standard Error | t value | p-value |
|---|---|---|---|---|
| Period-Wet | -0.0322 | 0.63306 | -0.051 | 0.96 |
| Sex-Male | 0.64104 | 0.45099 | 1.421 | 0.163 |
| Group-Coqueiro | -0.072 | 1.01245 | -0.071 | 0.944 |
| Group-Cow | -0.90477 | 1.35047 | -0.67 | 0.507 |
| Group-F Group | -0.9508 | 1.22953 | -0.773 | 0.444 |
| Group-House | -0.16544 | 0.88203 | -0.188 | 0.852 |

| | | | | |
|---|---|---|---|---|
| Group-Key | 0.47293 | 1.09268 | 0.433 | 0.668 |
| Group-Princess | -0.6155 | 0.87376 | -0.704 | 0.485 |
| Group-Road | -0.5351 | 1.12079 | -0.477 | 0.636 |
| Preservative-RNAlater | -0.24328 | 0.57327 | -0.424 | 0.674 |
| Age-Infant | 0.29425 | 0.96709 | 0.304 | 0.763 |
| Age-Juvenile | -0.3449 | 0.65467 | -0.527 | 0.601 |
| Age-Subadult | -0.01837 | 1.06814 | -0.017 | 0.986 |

**Table S16.** The results of a fixed-effects model that tested which parameter levels impacted the Simpson diversity of the likely resident fungi. No parameters were significant, and the overall model was not significant (p = 0.6564).

| Parameter-Level | Estimate | Standard Error | t value | p-value |
|---|---|---|---|---|
| Period-Wet | -0.08217 | 0.190141 | -0.432 | 0.668 |
| Sex-Male | 0.223941 | 0.135457 | 1.653 | 0.107 |
| Group-Coqueiro | 0.324016 | 0.30406 | 1.066 | 0.293 |
| Group-Cow | 0.497391 | 0.405616 | 1.226 | 0.228 |
| Group-F Group | 0.488094 | 0.369291 | 1.322 | 0.194 |
| Group-House | 0.354583 | 0.264919 | 1.338 | 0.189 |
| Group-Key | 0.488822 | 0.32819 | 1.489 | 0.145 |
| Group-Princess | 0.16538 | 0.262435 | 0.63 | 0.532 |
| Group-Road | -0.06512 | 0.336633 | -0.193 | 0.848 |
| Preservative-RNAlater | -0.05278 | 0.172184 | -0.307 | 0.761 |
| Age-Infant | -0.05765 | 0.290468 | -0.198 | 0.844 |
| Age-Juvenile | 0.007302 | 0.196633 | 0.037 | 0.971 |
| Age-Subadult | 0.195331 | 0.320817 | 0.609 | 0.546 |

**Table S17.** The results of a fixed-effects model that tested which parameter levels impacted the Pielou's evenness values of the likely resident fungi. No parameters were significant, and the overall model was not significant (p = 0.5152).

| Parameter-Level | Estimate | Standard Error | t value | p-value |
|---|---|---|---|---|
| Period-Wet | -0.21926 | 0.288171 | -0.761 | 0.4561 |
| Sex-Male | 0.328276 | 0.190091 | 1.727 | 0.1004 |
| Group-Coqueiro | -0.77209 | 0.439172 | -1.758 | 0.0948 |
| Group-Cow | -1.09279 | 0.540701 | -2.021 | 0.0576 |
| **Group-F Group** | **-1.29217** | **0.515993** | **-2.504** | **0.0215** |
| Group-House | -0.79637 | 0.3839 | -2.074 | 0.0519 |
| Group-Key | -0.81143 | 0.393321 | -2.063 | 0.053 |
| Group-Princess | -0.97832 | 0.473196 | -2.067 | 0.0526 |
| **Group-Road** | **-1.36899** | **0.648577** | **-2.111** | **0.0483** |

| | | | | |
|---|---|---|---|---|
| Preservative-RNAlater | -0.05694 | 0.237626 | -0.24 | 0.8132 |
| Age-Infant | 0.007235 | 0.303774 | 0.024 | 0.9812 |
| Age-Juvenile | 0.039761 | 0.246786 | 0.161 | 0.8737 |
| Age-Subadult | 0.253569 | 0.610282 | 0.415 | 0.6824 |

**Table S18.** The results of the ANCOM-BC, indicating which ASVs were differentially abundant between the sample collection periods. The drier period is the reference, with these values being associated with the wetter period.

| ASV | Taxon | LFC | Standard Error | p-value |
|---|---|---|---|---|
| 09747f28421a8d60ef07b64fdd7263b0 | *Nothophoma* sp. | -5.042 | 0.511 | 4.816E-05 |
| e462722cb5bc4b112adfc8babf18e384 | *Cladosporium* 1 | 3.679 | 0.371 | 4.816E-05 |
| 18aeb211ab36e34ea362709b418599cb | *Cladosporium* 2 | 1.813 | 0.295 | 4.163E-02 |
| 8c707734e4bd5cfc2f816d96dbd5cbaa | *Candida parapsilosis* | 1.431 | 0.379 | 1.337E-02 |
| b70aee864e2f9fdd5c1c3819d2b02e1a | *Penicillium sumatraense* | 1.590 | 0.295 | 3.438E-02 |

**Table S19.** The results of several Spearman correlation tests comparing the within sample diversity metrics of the plant portion of the marmoset diet to the corresponding metrics associated with the overall mycobiome, the likely transient fungi, and the likely resident fungi. None of the correlations were statistically significant.

| Comparison | Metric | p-value | rho |
|---|---|---|---|
| Overall Mycobiome | Shannon | 0.7661 | 0.04226074 |
| | Richness | 0.6078 | 0.07284217 |
| | Chao1 | 0.9287 | 0.01272091 |
| | Simpson | 0.4909 | -0.0976693 |
| | Pielou | 0.6147 | -0.0714591 |
| Likely Transient Fungi | Shannon | 0.889 | 0.01984266 |
| | Richness | 0.4442 | 0.1084232 |
| | Chao1 | 0.902 | 0.01741878 |
| | Simpson | 0.3738 | 0.1258992 |
| | Pielou | 0.3341 | -0.1547308 |
| Likely Resident Fungi | Shannon | 0.3688 | 0.1272089 |
| | Richness | 0.4424 | 0.1088634 |
| | Chao1 | 0.2109 | 0.176059 |
| | Simpson | 0.8125 | -0.033703 |
| | Pielou | 0.7612 | 0.05497862 |

**Table S20.** The results of several Spearman correlation tests comparing the within sample diversity metrics of the arthropod portion of the marmoset diet to the corresponding metrics associated with the overall mycobiome, the likely transient fungi, and the likely resident fungi. Significant correlations are bolded.

| Comparison | Metric | p-value | rho |
|---|---|---|---|
| Overall Mycobiome | Shannon | 0.9533 | 0.00832426 |
| | **Richness** | **0.03906** | **0.2870812** |
| | Chao1 | 0.1289 | 0.2133037 |
| | Simpson | 0.8424 | -0.0282598 |
| | Pielou | 0.1912 | -0.1841583 |
| Likely Transient Fungi | Shannon | 0.7145 | -0.0519598 |
| | **Richness** | **0.03534** | **0.292522** |
| | Chao1 | 0.09028 | 0.2373073 |
| | Simpson | 0.2402 | -0.1657772 |
| | Pielou | 0.8673 | 0.02692224 |
| Likely Resident Fungi | Shannon | 0.5722 | 0.08015305 |
| | Richness | 0.1498 | 0.202564 |
| | Chao1 | 0.3954 | 0.1203612 |
| | **Simpson** | **0.03196** | **-0.2978917** |
| | Pielou | 0.8591 | 0.03213546 |

**Table S21.** The results of a CCREPE analysis investigating interactions between 12 fungal ASVs and 44 arthropod ASVs. Each row represents a pairwise interaction, with columns showing the fungal ASV, arthropod ASV, p-value, q-value, sim score, and z stat. Table is included as a separate file titled SupplementalTable21.xlsx

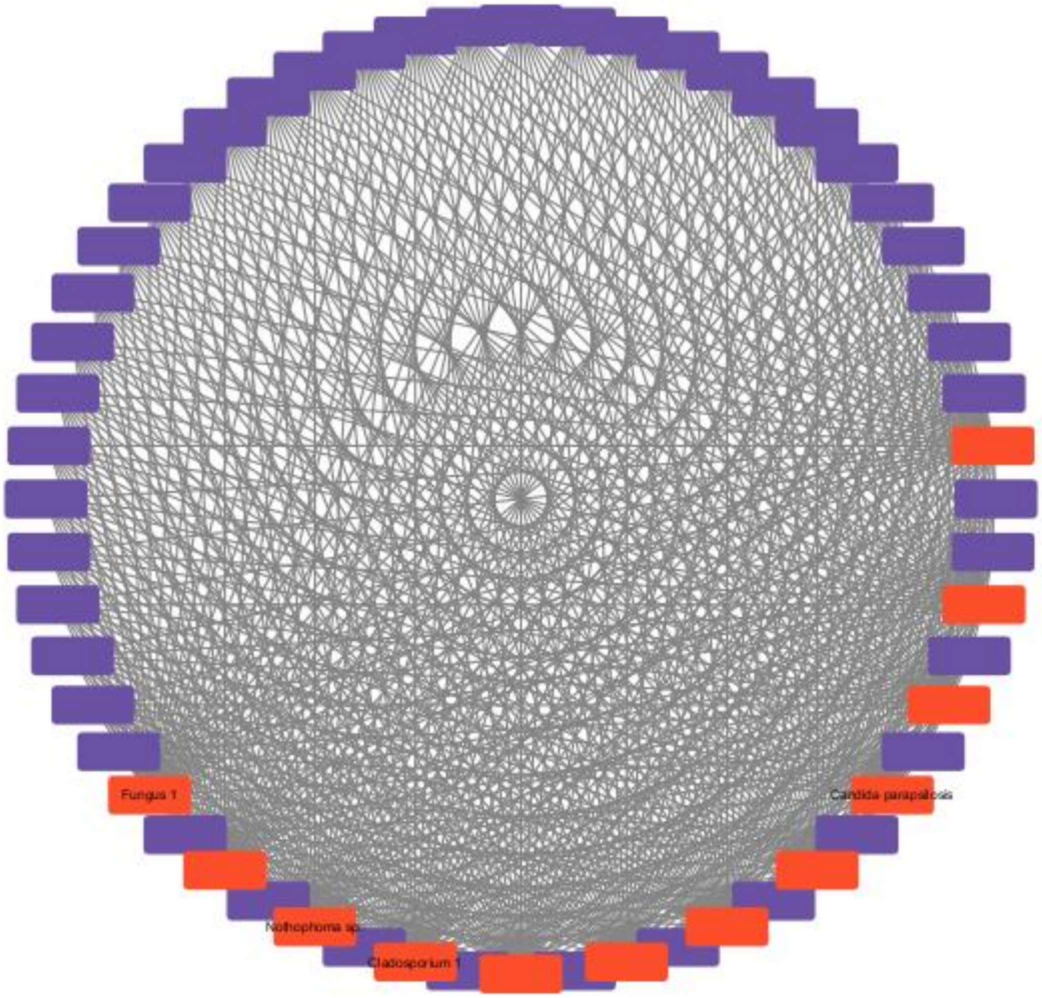

**Figure S8.** Network of interactions between fungal ASVs (red) and arthropod ASVs (purple). No interaction was significant (q-value < 0.05 and absolute sim score > 0.6; Table S21). Fungal ASVs that were identified by ANCOM-BC are labeled.

**Table S22.** The results of a CCREPE analysis investigating interactions between 12 fungal ASVs and 266 plant ASVs. Each row represents a pairwise interaction, with columns showing the fungal ASV, arthropod ASV, p-value, q-value, sim score, and z stat. The significant relationship was bolded and moved to the top of the table. Table is included as a separate file titled SupplementalTable22.xlsx

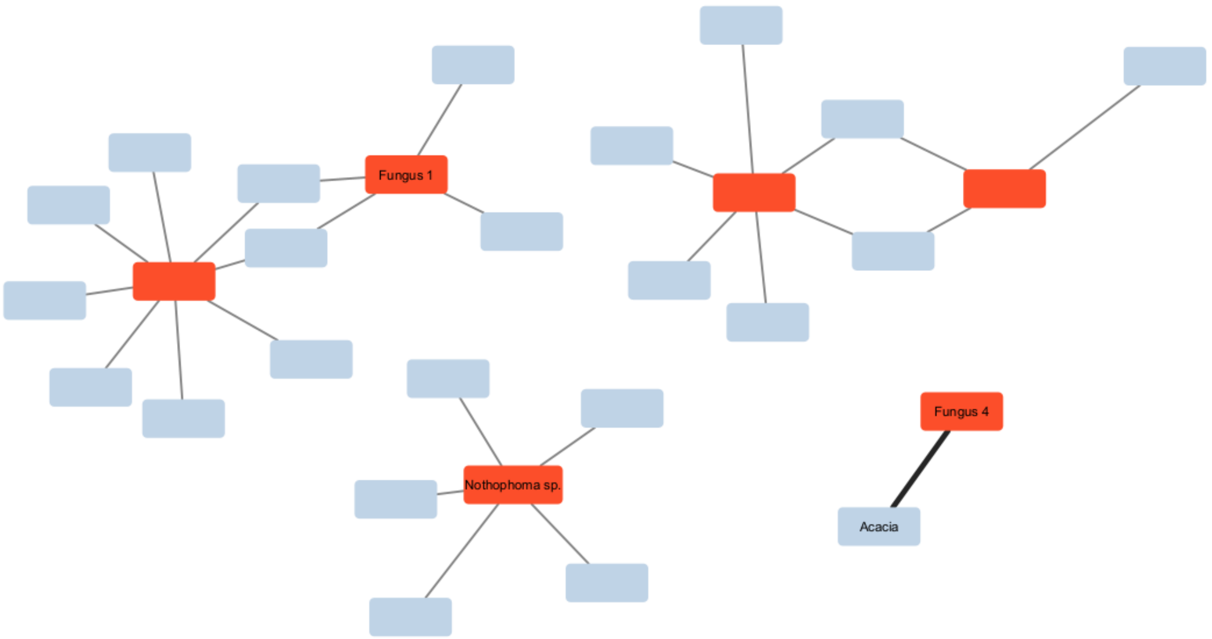

**Figure S9.** Network of interactions between fungal ASVs (red) and plant ASVs (gray). Only ASVs involved in interactions with q-values below 1 were mapped. Significant interactions are indicated by thick bolded edges (Table S22). ASVs that were identified as significant by ANCOM-BC and/or CCREPE are labeled.

**Table S23.** Results of likelihood ratio tests comparing full mixed-effects models to null models missing the sex parameter to determine whether it significantly relates to within sample diversity metrics of arthropod portion of the arthropod diet. None of the models were significant.

| Diversity Metric | Model | logLik | deviance | Chisq | Df | Pr(>Chisq) |
|---|---|---|---|---|---|---|
| Shannon | Full Model | -43.8 | 87.6 | 0.8283 | 1 | 0.3628 |
| | Null Sex Model | -44.214 | 88.429 | | | |
| Richness | Full Model | -219.32 | 438.64 | 0.1003 | 1 | 0.7515 |
| | Null Sex Model | -219.7 | 438.74 | | | |
| Chao1 | Full Model | -250.95 | 501.89 | 2.0722 | 1 | 0.15 |
| | Null Sex Model | -251.98 | 503.97 | | | |
| Simpson | Full Model | 16.279 | -32.557 | 0.52 | 1 | 0.4709 |
| | Null Sex Model | 16.019 | -32.037 | | | |

**Table S24.** The results of a fixed-effects model that tested which parameter levels related to the Pielou's evenness of the arthropod portion of the marmoset diet. None of the parameters were significant.

| Parameter-Level | Estimate | Standard Error | t-value | p-value |
|---|---|---|---|---|
| (Intercept) | 0.475 | 0.138 | 3.431 | 0.001 |
| Period-Wet | -0.047 | 0.100 | -0.473 | 0.639 |
| Sex-Male | 0.005 | 0.071 | 0.075 | 0.940 |
| Group-Coqueiro | 0.092 | 0.159 | 0.575 | 0.569 |
| Group-Cow | 0.048 | 0.213 | 0.226 | 0.823 |
| Group-F group | 0.306 | 0.193 | 1.580 | 0.122 |
| Group-House | 0.132 | 0.139 | 0.952 | 0.347 |
| Group-Key | 0.080 | 0.172 | 0.468 | 0.643 |
| Group-Princess | 0.149 | 0.138 | 1.080 | 0.287 |
| Group-Road | -0.110 | 0.176 | -0.624 | 0.537 |
| Preservative-RNAlater | -0.021 | 0.090 | -0.231 | 0.818 |
| Age-Infant | -0.158 | 0.152 | -1.036 | 0.307 |
| Age-Juvenile | 0.100 | 0.103 | 0.966 | 0.340 |
| Age-Subadult | 0.127 | 0.168 | 0.757 | 0.454 |

**Table S25.** The results of a fixed-effects model that tested which parameter levels related to the Shannon diversity of the plant portion of the marmoset diet. None of the parameters were significant.

| Parameter-Level | Estimate | Standard Error | t-value | p-value |
|---|---|---|---|---|
| (Intercept) | 1.867 | 0.656 | 2.847 | 0.007 |
| Period-Wet | -0.102 | 0.472 | -0.216 | 0.830 |
| Sex-Male | -0.461 | 0.337 | -1.369 | 0.179 |
| Group-Coqueiro | 0.458 | 0.756 | 0.606 | 0.548 |
| Group-Cow | 1.385 | 1.008 | 1.375 | 0.177 |
| Group-F group | -0.135 | 0.918 | -0.147 | 0.884 |
| Group-House | 0.128 | 0.658 | 0.195 | 0.847 |
| Group-Key | 0.266 | 0.815 | 0.326 | 0.746 |
| Group-Princess | -0.032 | 0.652 | -0.049 | 0.961 |
| Group-Road | 0.336 | 0.836 | 0.402 | 0.690 |
| Preservative-RNAlater | 0.486 | 0.428 | 1.135 | 0.263 |
| Age-Infant | 0.448 | 0.722 | 0.621 | 0.539 |
| Age-Juvenile | 0.040 | 0.489 | 0.083 | 0.935 |
| Age-Subadult | -0.108 | 0.797 | -0.136 | 0.893 |

**Table S26.** Results of likelihood ratio tests comparing full mixed-effects models to null models missing the sex parameter to determine whether it has significant impacts on within sample diversity metrics of arthropod portion of the arthropod diet. None of the models were significant.

| Diversity Metric | Model | logLik | deviance | Chisq | Df | Pr(>Chisq) |
|---|---|---|---|---|---|---|
| Richness | Full Model | -331.6 | 663.2 | | | |
| | Null Sex Model | -329.93 | 659.86 | 3.3405 | 1 | 0.06759 |
| Chao1 | Full Model | -344.02 | 688.05 | 0.6554 | 1 | 0.4182 |
| | Null Sex Model | -344.35 | 688.7 | | | |

**Table S27.** The results of a fixed-effects model that tested which parameter levels related to the Simpson diversity of the plant portion of the marmoset diet. None of the parameters of interest were significant.

| Parameter-Level | Estimate | Standard Error | t-value | p-value |
|---|---|---|---|---|
| (Intercept) | 0.516 | 0.176 | 2.941 | 0.006 |
| Period-Wet | -0.018 | 0.126 | -0.144 | 0.886 |
| Sex-Male | -0.148 | 0.090 | -1.648 | 0.108 |
| Group-Coqueiro | 0.162 | 0.202 | 0.799 | 0.429 |
| Group-Cow | 0.372 | 0.270 | 1.379 | 0.176 |
| Group-F group | -0.049 | 0.246 | -0.201 | 0.842 |
| Group-House | 0.056 | 0.176 | 0.318 | 0.752 |
| Group-Key | 0.093 | 0.218 | 0.425 | 0.673 |
| Group-Princess | -0.001 | 0.175 | -0.007 | 0.995 |
| Group-Road | 0.133 | 0.224 | 0.592 | 0.557 |
| Preservative-RNAlater | 0.163 | 0.115 | 1.426 | 0.162 |
| Age-Infant | 0.081 | 0.193 | 0.418 | 0.678 |
| AgeJ-uvenile | 0.007 | 0.131 | 0.056 | 0.956 |
| Age-Subadult | -0.099 | 0.213 | -0.465 | 0.645 |

**Table S28.** The results of a fixed-effects model that tested which parameter levels related to the Pielou's evenness of the plant portion of the marmoset diet. None of the parameters of interest were significant.

| Parameter-Level | Estimate | Standard Error | t-value | p-value |
|---|---|---|---|---|
| (Intercept) | 0.383 | 0.197 | 1.947 | 0.059 |
| Period-Wet | -0.069 | 0.142 | -0.488 | 0.628 |
| Sex-Male | -0.169 | 0.101 | -1.675 | 0.102 |
| Group-Coqueiro | 0.222 | 0.227 | 0.978 | 0.334 |
| Group-Cow | 0.451 | 0.303 | 1.490 | 0.144 |
| Group-F group | -0.080 | 0.275 | -0.290 | 0.774 |

| | | | | |
|---|---|---|---|---|
| Group-House | -0.016 | 0.198 | -0.082 | 0.935 |
| Group-Key | 0.108 | 0.245 | 0.440 | 0.662 |
| Group-Princess | 0.032 | 0.196 | 0.163 | 0.872 |
| Group-Road | 0.141 | 0.251 | 0.560 | 0.579 |
| Preservative-RNAlater | 0.300 | 0.128 | 2.336 | 0.025 |
| Age-Infant | 0.012 | 0.217 | 0.053 | 0.958 |
| AgeJ-uvenile | -0.024 | 0.147 | -0.167 | 0.869 |
| Age-Subadult | -0.153 | 0.239 | -0.638 | 0.527 |

**Table S29.** The results of a PERMANOVA using marginal sums of squares looking at the effects of various parameters on the Bray-Curtis distances between the compositions of arthropod portion of the diet of common marmoset samples. Significant parameters are bolded.

| Parameter | Degrees of Freedom | Sum of Squares | R2 | Pseudo-F | p-value |
|---|---|---|---|---|---|
| Age | 3 | 1.3751 | 0.0563 | 0.9684 | 0.638 |
| Sex | 1 | 0.4147 | 0.1698 | 0.8762 | 0.828 |
| **Period** | **1** | **0.6573** | **0.2691** | **1.3887** | **0.01** |
| Group | 7 | 3.5395 | 0.1449 | 1.0683 | 0.111 |
| Preservative | 1 | 0.4699 | 0.01924 | 0.9929 | 0.493 |
| Residuals | 38 | 17.9863 | 0.73633 | | |

**Table S30.** The results of a PERMANOVA using marginal sums of squares looking at the effects of various parameters on the Jaccard distances between the compositions of arthropod portion of the diet of common marmoset samples. Significant parameters are bolded.

| Parameter | Degrees of Freedom | Sum of Squares | R2 | Pseudo-F | p-value |
|---|---|---|---|---|---|
| Age | 3 | 1.2498 | 0.05571 | 0.963 | 0.605 |
| Sex | 1 | 0.4041 | 0.01801 | 0.9341 | 0.615 |
| **Period** | **1** | **0.6277** | **0.02798** | **1.4509** | **0.025** |
| Group | 7 | 3.1573 | 0.14073 | 1.0426 | 0.261 |
| Preservative | 1 | 0.5067 | 0.02259 | 1.1713 | 0.143 |
| Residuals | 38 | 16.4398 | 0.73276 | | |

**Table S31.** The results of a PERMANOVA using marginal sums of squares looking at the effects of various parameters on the Bray-Curtis distances between the compositions of plant portion of the diet of common marmoset samples. Significant parameters are bolded.

| Parameter | Degrees of Freedom | Sum of Squares | R2 | Pseudo-F | p-value |
|---|---|---|---|---|---|
| Age | 3 | 0.8613 | 0.04715 | 0.8472 | 0.64 |
| Sex | 1 | 0.4162 | 0.02278 | 1.2282 | 0.256 |
| Period | 1 | 0.3173 | 0.01737 | 0.9363 | 0.443 |

| | | | | | |
|---|---|---|---|---|---|
| Group | 7 | 2.5277 | 0.13837 | 1.0656 | 0.347 |
| **Preservative** | **1** | **1.0496** | **0.05745** | **3.0971** | **0.012** |
| Residuals | 38 | 12.8778 | 0.70493 | | |

**Table S32.** The results of a PERMANOVA using marginal sums of squares looking at the effects of various parameters on the Jaccard distances between the compositions of plant portion of the diet of common marmoset samples. Significant parameters are bolded.

| Parameter | Degrees of Freedom | Sum of Squares | R2 | Pseudo-F | p-value |
|---|---|---|---|---|---|
| Age | 3 | 0.8892 | 0.04768 | 0.8498 | 0.799 |
| Sex | 1 | 0.3438 | 0.01844 | 0.9858 | 0.385 |
| Period | 1 | 0.2833 | 0.01519 | 0.8123 | 0.68 |
| Group | 7 | 2.6558 | 0.14241 | 1.0878 | 0.214 |
| **Preservative** | **1** | **0.896** | **0.04804** | **2.5689** | **0.005** |
| Residuals | 38 | 13.2534 | 0.71066 | | |

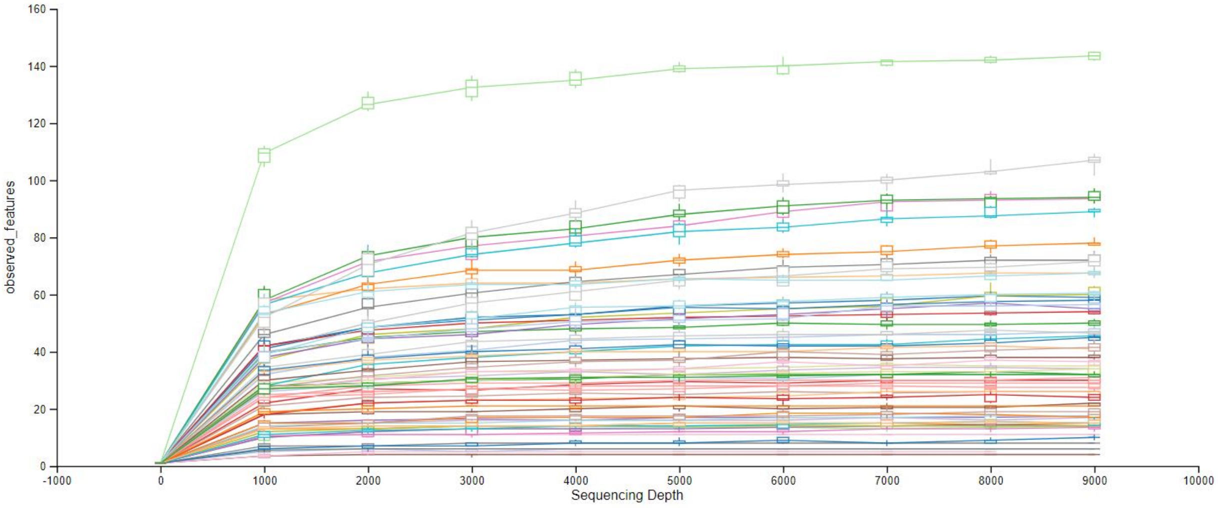

**Figure S10.** The rarefaction plot generated by Qiime2. Based on this plot, a depth of 5,000 reads was selected for rarefaction.

**Table S33.** A list of the samples that had fewer than 5,000 reads and their relevant metadata.

| Sample | Group | Age | Sex | Preservative | Reproductive | Period | Time |
|---|---|---|---|---|---|---|---|
| CJAC18 | House | Unknown | Unknown | RNAlater | Unknown | Dry | Dry2015 |
| CJAC21 | Algaroba | Unknown | Unknown | RNAlater | Unknown | Dry | Dry2015 |
| CJAC22 | House | Adult | Female | RNAlater | ND | Dry | Dry2015 |
| CJAC25 | House | Adult | Male | RNAlater | NPMale | Dry | Dry2015 |

| | | | | | | | |
|---|---|---|---|---|---|---|---|
| CJAC36 | Cow | Adult | Male | RNAlater | NPMale | Wet | Wet2015 |
| CJAC37 | Cow | Adult | Male | RNAlater | NPMale | Wet | Wet2015 |
| CJAC43 | Cow | Adult | Female | RNAlater | Unknown | Wet | Wet2015 |

**Table S34.** The MDS1 and MDS2 values from the non-metric multidimensional scaling plot including all samples. Samples that disproportionately impacted the plot are bolded.

| Sample | MDS1 | MDS2 |
|---|---|---|
| CJAC10 | -3.067 | -15.654 |
| CJAC101 | -3.064 | -15.714 |
| CJAC102 | -3.068 | -15.646 |
| CJAC104 | -3.010 | -15.587 |
| CJAC105 | -3.075 | -15.612 |
| CJAC107 | -3.545 | -15.952 |
| CJAC11 | -3.069 | -15.605 |
| CJAC114 | -3.052 | -15.606 |
| CJAC115 | -3.077 | -15.651 |
| CJAC116 | -3.118 | -15.689 |
| CJAC117 | -3.010 | -15.650 |
| CJAC118 | -3.138 | -15.625 |
| CJAC12 | -3.088 | -15.650 |
| CJAC125 | -3.055 | -15.618 |
| CJAC126 | -3.076 | -15.615 |
| CJAC127 | -3.055 | -15.637 |
| CJAC129 | -3.078 | -15.609 |
| CJAC13 | -3.046 | -15.625 |
| CJAC132 | -3.085 | -15.575 |
| CJAC134 | -3.053 | -15.723 |
| CJAC135 | -3.103 | -15.574 |
| CJAC138 | -3.110 | -15.516 |
| CJAC14 | -3.048 | -15.615 |
| CJAC19 | -3.080 | -15.665 |
| CJAC2 | -3.031 | -15.562 |
| CJAC20 | -2.926 | -15.527 |
| CJAC24 | -3.056 | -15.601 |
| CJAC32 | -2.912 | -15.381 |
| CJAC35 | -3.018 | -15.691 |
| CJAC4 | -3.142 | -15.635 |
| CJAC48 | -3.042 | -15.607 |
| CJAC5 | -3.036 | -15.650 |
| CJAC55 | -3.120 | -15.618 |
| CJAC56 | -3.071 | -15.614 |

| | | |
|---|---|---|
| CJAC6 | -3.077 | -15.613 |
| CJAC63 | -3.041 | -15.638 |
| CJAC64 | -3.120 | -15.560 |
| CJAC66 | -3.087 | -15.619 |
| CJAC7 | -3.091 | -15.621 |
| CJAC71 | -3.127 | -15.568 |
| CJAC74 | -3.013 | -15.637 |
| CJAC76 | -3.130 | -15.597 |
| **CJAC79** | **2789.444** | **378.885** |
| CJAC8 | -3.111 | -15.631 |
| CJAC83 | -3.117 | -15.630 |
| CJAC84 | -3.062 | -15.634 |
| **CJAC85** | **-2635.343** | **401.954** |
| CJAC86 | -3.101 | -15.599 |
| CJAC87 | -3.142 | -15.581 |
| CJAC89 | -3.028 | -15.645 |
| CJAC94 | -3.068 | -15.561 |
| CJAC95 | -3.259 | -15.406 |

**Table S35.** A list of the samples that had a disproportionate effect on the non-metric multidimensional scaling plot of the Bray-Curtis distance plot and their relevant metadata.

| Sample | Group | Age | Sex | Preservative | Reproductive | Period | Time |
|---|---|---|---|---|---|---|---|
| CJAC79 | Princess | Adult | Female | RNAlater | Pregnant | Wet | Wet2015 |
| CJAC85 | Princess | Adult | Male | RNAlater | Non-pregnant Male | Wet | Wet2015 |

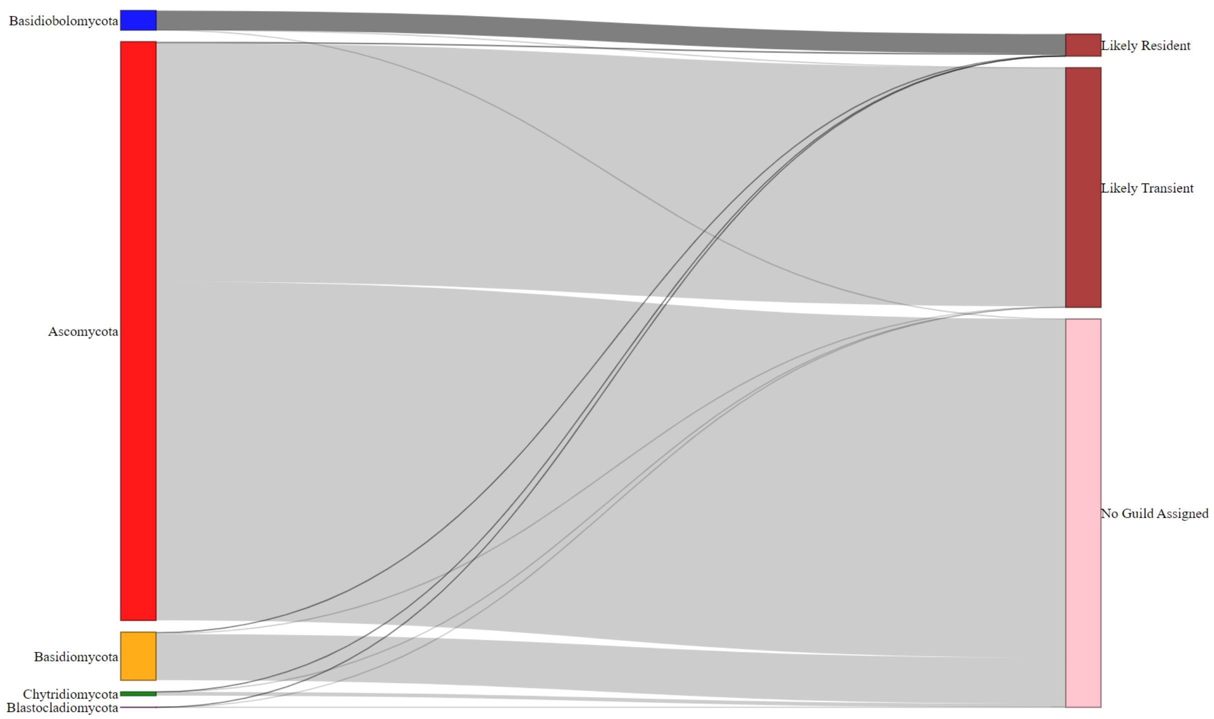

**Figure S11.** A Sankey plot showing how reads from the detected phyla contribute to the fungal categories of interest. Note, ASVs that were not assigned to any phylum are excluded from this figure.

**Table S36.** A list of samples that did not have information about the sex of the sampled individual and were removed from linear model and PERMANOVA analyses.

| Sample | Group | Age | Sex | Preservative | Reproductive | Period | Time |
|---|---|---|---|---|---|---|---|
| CJAC15 | Algaroba | Unknown | Unknown | RNAlater | Unknown | Dry | Dry2015 |
| CJAC16 | Algaroba | Infant | Unknown | RNAlater | Not pregnant | Dry | Dry2015 |
| CJAC23 | House | Unknown | Unknown | RNAlater | Unknown | Dry | Dry2015 |
| CJAC34 | Cow | Unknown | Unknown | RNAlater | Unknown | Wet | Wet2015 |
| CJAC42 | Cow | Unknown | Unknown | RNAlater | Unknown | Wet | Wet2015 |
| CJAC69 | Key | Unknown | Unknown | RNAlater | Unknown | Wet | Wet2015 |
| CJAC72 | Princess | Unknown | Unknown | RNAlater | Unknown | Wet | Wet2015 |
| CJAC96 | Princess | Adult | Unknown | Ethanol | Unknown | Dry | Dry2016 |

**Table S37.** A list of diversity metrics calculated for the mycobiome, transient mycobiome, and resident mycobiome with information about which modeling approach was used to analyze it.

| Diversity Metric | Overall Mycobiome | Transient Mycobiome | Resident Mycobiome |
|---|---|---|---|
| Shannon | Mixed-Effect | Fixed-Effect | Fixed-Effect |
| Richness | Mixed-Effect | Mixed-Effect | Fixed-Effect |
| Chao1 | Mixed-Effect | Mixed-Effect | Fixed-Effect |
| Simpson | Fixed-Effect | Fixed-Effect | Fixed-Effect |
| Pielou's Evenness | Fixed-Effect | Fixed-Effect | Fixed-Effect |
